# Mutational, transcriptional and viral shedding dynamics of the marine turtle fibropapillomatosis tumor epizootic

**DOI:** 10.1101/2020.02.04.932632

**Authors:** Kelsey Yetsko, Jessica Farrell, Maximilian R. Stammnitz, Liam Whitmore, Jenny Whilde, Catherine B. Eastman, Devon Rollinson Ramia, Rachel Thomas, Aleksandar Krstic, Paul Linser, Simon Creer, Gary Carvalho, Brooke Burkhalter, Elizabeth P. Murchison, Christine Schnitzler, David J. Duffy

## Abstract

Sea turtle populations are directly and indirectly under threat from a range of anthropogenic processes. Perhaps the most visibly apparent of these is the disfiguring tumor disease epizootic (animal epidemic) known as fibropapillomatosis. Fibropapillomatosis continues to spread geographically, with prevalence of the disease also growing at a number of affected sites globally. Environmental exposures seem key to inducing tumor development, possibly through weakening host immune systems to the point of enabling pathogen-induced tumor formation. However, we do not yet understand the precise molecular and mutational events driving fibropapillomatosis tumor formation and progression. Similarly, many open questions remain about the role of the herpesvirus (chelonid herpesvirus 5, ChHV5) associated with the disease as a potential co-trigger, and whether its occurrence within tumors is causative or opportunistic. Without improved understanding of the basic biology of this disease epizootic, treatment, containment and mitigation options are severely hampered.

To address fundamental questions relating to the oncogenic signaling, mutational spectrum, viral load, viral transcriptional status (lytic or latent) and spread, we employed transcriptomic profiling, whole genome sequencing, immunohistochemistry and environmental (e)DNA-based monitoring of viral shedding. In particular we focused on the mutational landscape of tumors and assessing the transcriptional similarity of external (skin) and internal (visceral organs) tumors, and the oncogenic signaling events driving early stage tumor growth and post-surgical tumor regrowth. These analyses revealed that internal fibropapillomatosis tumors are molecularly distinct from the more common external tumors. However, our molecular analyses also revealed that there are a small number of conserved potentially therapeutically targetable molecular vulnerabilities in common between internal and external tumors, such as the MAPK, Wnt, TGFβ and TNF oncogenic signaling pathways. We also determined that the tumor genomes can harbor copy number gains, indicating potentially viral-independent oncogenic processes. Genes within such mutated genomic regions have known roles in human skin cancer, including MAPK-associated genes. Turtles attempt to mount an immune response, but in some animals this appears to be insufficient to prevent tumor development and growth. ChHV5 was transcriptionally latent in all tumor stages sequenced, including early stage and recurrent tumors. We also revealed that the tumors themselves are the primary source of viral shedding into the marine environment and, if they are surgically removed, the level of ChHV5 in the water column drops.

Together, these results offer an improved understanding of fibropapillomatosis tumorigenesis and provide insights into the origins, therapeutic treatment, and appropriate quarantine responses for this wildlife epizootic. Furthermore, they provide insights into human pathogen-induced cancers, particularly mechanisms which are difficult to study in the human and terrestrial context, such as time-course quantification-based monitoring of viral shedding.

## Introduction

Sea turtle fibropapillomatosis (FP) is potentially a canary in the coalmine, indicating that continued human-induced environmental damage may be an alternative route by which oncogenicity is conferred on normally well-tolerated viruses. This is particularly worrying as long-lived reptiles have normally robust anti-cancer defenses^1, 2^, and as there are already a range of human viruses known to be capable of inducing tumor formation when the host immune system is compromised^3, 4^. In addition to improving wildlife conservation medicine, fibropapillomatosis precision oncology can reveal the precise mechanisms through with environmental triggers, viral dynamics and host cell transformation can rapidly induce novel cancer incidence on an epidemic scale, thereby simultaneously informing human cancer research^2, 5^.

Fibropapillomatosis (Fig. 1A) is a tumor disease event of epizootic (animal epidemic) proportions, affecting wild populations of endangered marine turtles circum-globally^6–9^. Fibropapillomatosis leads to sea turtle fatalities by direct (affecting internal organs) and indirect means (blocking vision and feeding or restricting movement and immune functioning)^7, 10^. This sea turtle tumor epizootic is particularly striking given that long-lived turtle species (over 80 years) tend to have more robust anti-cancer defenses than humans^1^. Green sea turtles (*Chelonia mydas*) are most affected by fibropapillomatosis, providing a clear species to focus on for experimental studies, but the disease also occurs in all other marine turtle species^11–14^. Sea turtle fibropapillomatosis continues to spread geographically^7, 10^ throughout equatorial and subequatorial regions^7, 10, 11, 15, 16^. Fibropapillomatosis has been reported in every major ocean basin in which green turtles are found, particularly in near-shore habitats^13, 17^(www.cabi.org/isc/datasheet/82638). In addition to spreading globally, fibropapillomatosis rates continue to increase in many affected sites, posing serious conservation challenges. Currently, of all green sea turtles stranding, over 40% in Florida, 30% in Hawaii, 35% in Texas, 35% in northeastern Brazil, 34% at Príncipe Island in the Gulf of Guinea and 50% in Puerto Rico are FP-afflicted ^10, 18–22^. Many of these sites have seen a rapid increase in disease incidence over recent years, for instance 13.3% in 2005 compared with 42% in 2016 in Florida, 13.2% in 2012 compared with 35.3% in 2015 in northeastern Brazil, and 0% in 2009 compared with 35.2% in 2018 in Texas (the occurrence of fibropapillomatosis in Texas was first reported in 2010 at a rate of 0.6%) ^10, 18–21^.

**Figure 1.**
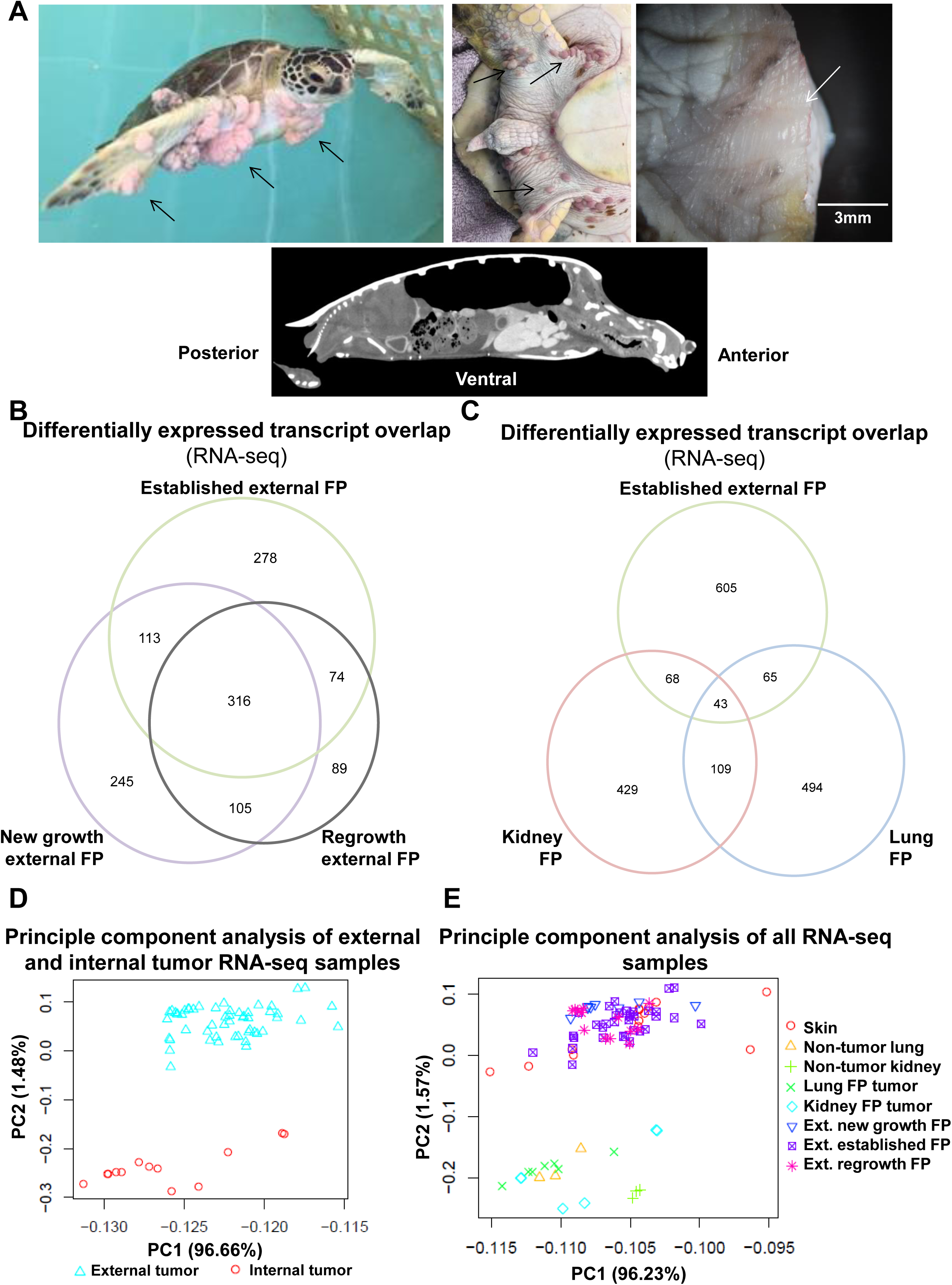
Fibropapillomatosis tumors and differential transcript expression. **A)** Top, left: Fibropapillomatosis-afflicted green sea turtles (*Chelonia mydas*) in one of the hospital’s seawater tanks, awaiting tumor removal surgery. Established tumors are visible as large pinkish outgrowths. Top, middle: Numerous new growth tumors occurring around the ventral tail and rear flipper area of patient 25-2018-Cm ‘Lilac’. Top, right: Post-surgical regrowth tumor imaged after surgical resection. Re-growing tumor is pinkish tissue surrounded by paler non-tumored skin.Bottom: A computed tomography (CT) scan of fibropapillomatosis-afflicted *C. mydas*. CT is one of the approaches used for diagnosing internal tumors. In all images arrows indicate selected examples of tumors. **B, C)** Overlap of transcripts significantly differentially expressed (DE) (as called by DESeq2) in fibropapillomatosis from the RNA-seq data. Area-proportional Venn diagrams were generated using BioVenn (http://www.biovenn.nl/)^97^). Transcripts were considered significant if passing the following cut-offs: adjusted p-value of < 0.05 and log_2_ fold change of > 2 or < −2. **B)** Overlap of DEs from the following comparisons: established external FP, new growth external FP, and regrowth external FP, when all are compared to healthy skin for differential expression analysis. **C)** Overlap of DEs from the following comparisons: established external FP, kidney FP, and lung FP, each compared to their non-tumored tissue sources for differential expression analysis (healthy skin, healthy kidney, and healthy lung, respectively). **D)** Principle component analysis (PCA) of all internal tumor samples compared to external tumor samples, RNA-seq. **E)** PCA of all samples, includes all tumor and non-tumor samples, RNA-seq.

A chelonian-specific herpesvirus (chelonid herpesvirus 5, ChHV5), has been implicated in driving this disease epizootic, although Koch’s postulates to confirm its causative role have yet to be definitively confirmed ^7, 12, 23–26^. Even so, ChHV5 infection alone is not sufficient to induce FP tumor growth. ChHV5 is globally distributed, including being present in sea turtle populations in which fibropapillomatosis tumors have never been reported^27^. An environmental co-trigger(s) seems to be the required key to both the development of fibropapillomatosis tumors and the geographic spread of the disease, with the environmental trigger correlating with anthropogenic activity^7, 26, 28, 29^. Much uncertainty remains about the specific environmental trigger(s) and how they interact with ChHV5 and the host immune-system to give rise to FP tumorigenesis^7^. It is highly likely that environmental exposures induce immunosuppression in marine turtles enabling viral load to increase to the point of crossing an oncogenic threshold, similar to a number of human virally-induced cancers ^3, 7, 26, 30^. There is a paucity of knowledge concerning the molecular signaling events underpinning fibropapillomatosis tumor initiation, development and growth, with even less known about the viral and host transcriptional and mutational landscape driving FP tumorigenesis. Additionally, the precise relationship between internal and the more common external fibropapillomatosis remains to be elucidated.

Similarly, open questions remain regarding whether ChHV5 is lytic or latent in FP tumors and the occurrence of viral shedding and transmission^7, 26, 31–34^. Novel environmental DNA (eDNA) approaches can help resolve the dynamics of viral shedding. Environmental DNA is a forensics approach to the extraction and identification of organismal DNA fragments (genetic material) released into the environment and is a rapidly advancing, non-invasive approach capable of improving endangered species detection and early pathogen detection^35–43^. Environmental samples can be analyzed for micro- and macro-organisms by several eDNA methods including metabarcoding and species-specific quantitative PCR (qPCR)^38, 40, 44^. The development of a rapid and high throughput sampling scheme to detect virus shedding into the marine environment would benefit pathogen surveillance efforts, and consequently the performance of wildlife health status monitoring could be improved^31, 45^. Here we applied qPCR and whole genome sequencing/shotgun sequencing (“unbiased”, nonbarcoded) eDNA approaches to quantify ChHV5 viral shedding from rehabilitating patients. Such novel approaches to environmental viral shedding are a particularly beneficial feature of aquatic models of virally-induced tumors, such as fibropapillomatosis, and will enable greatly improved understanding of the dynamic relationship between viral load and viral shedding, not readily measurable in terrestrial species.

While advances in our understanding of the fibropapillomatosis tumor disease epizootic in sea turtles continue to be made^7, 26, 27, 30, 46–53^, many questions remain unanswered in relation to this enigmatic disease. There is virtually no molecular information about the relationship (e.g. primary/metastatic) between the numerous tumors (which can range from tens to hundreds of tumors per individual) arising on an individual. Similarly, the drivers of early stage, internal and post-surgical regrowth tumors remain to be elucidated. Determining the contribution of each facet of this multifactorial disease will be key to combatting this anthropogenic-implicated disease epidemic both at the level of individual clinical treatment and for population level management/mitigation strategies ^2, 30^. The relative contributions of ChHV5, environmental trigger(s), immune suppression, host genome mutation and altered gene expression to fibropapillomatosis oncogenesis have never been determined. Indeed, it is not known whether any mutational events occur within fibropapillomatosis tumor genomes. Here we applied a combination of extensive transcriptomics, genomics, precision oncology^30, 54, 55^, immunohistochemistry and eDNA-based viral shedding monitoring to determine the host and viral molecular events underpinning fibropapillomatosis tumorigenesis, growth, post-surgical recurrence and transmission.

## Results

### Fibropapillomatosis tumors oncogenic signaling networks, immune response and therapeutic vulnerabilities

To determine the molecular events governing FP tumor growth, and whether different oncogenic signaling networks drive varying tumor types in different tissues we conducted transcriptomics of fibropapillomatosis tumors and patient matched non-tumor tissue (90 RNA-seq samples, Supplemental Table 1). Differentially expressed genes were then analyzed at the gene, pathway, network, and systems level. When the mRNA transcripts differentially expressed between external new growth, external established, and external post-surgical regrowth (recurring) fibropapillomatosis tumors were compared, there was a high degree of overlap (Fig. 1B). This suggests that the molecular events driving fibropapillomatosis tumor formation, growth and regrowth are broadly similar.

Next we compared the mRNA transcripts differentially expressed (DE) between established external tumors and those of internal visceral tumors, lung FP and kidney FP (Fig. 1C, Supplemental Table 2). In contrast to the three types of external tumors (new growth, established, re-growth, Fig. 1B), the internal visceral tumor DE transcripts were extremely divergent to the external tumor DE transcripts (Fig. 1C). Furthermore, lung and kidney tumors also showed minimal overlap with each other (Fig. 1C) suggesting different oncogenic signals are driving internal FP compared with external FP tumors, and also kidney FP tumors compared with lung FP tumors. The diverging transcriptional profiles of internal and external tumors can also be seen at the whole transcriptome level (Fig. 1D,E).

A large number of genes were differentially expressed between all tumor samples and all non-tumor samples (Fig. 2A). Highly upregulated genes included those associated with oncogenic signaling, such as CTHRC1 (Wnt and adenocarcinoma signaling) and CRABP2 (retinoic acid signaling), while highly downregulated where associated with skeletal muscle (AACTA1, MYL6B and KLHL41, Fig. 2A). To better understand the biological processes involved in each tumor type we examined the top 20 Gene Ontology (GO) terms (called by IPA, ranked by p-value) for each set of differentially expressed transcripts (Fig. 2B,C and Supplemental Fig.1A-C). Eighteen of the top 20 GO terms associated with established external FP tumors were cancer or neoplasia-associated terms, with ‘activation of skin tumors’ and ‘melanoma’ associated terms featuring prominently (Fig. 2B). The two non-cancer GO terms, ‘formation of muscle’ and ‘muscle contraction’, were inhibited, likely indicating an absence of muscle in FP tumors compared with non-tumored skin controls. For new growth external tumors, 14 of the top 20 GO terms were cancer-associated (Supplemental Fig.1A). These 14 GO terms were not as highly activated as in established tumors, likely due to the early stage at which these tumors were sequenced, and the paucity of early stage (sequenced human) tumors in the IPA knowledgebase compared with established tumors (the stage more frequently diagnosed and biopsied/sequenced in human samples). In terms of level of activation, the GO terms of regrowth external tumors were intermediate between new growth and established FP. Eleven of the top 20 regrowth GO terms were cancer-related, again with a number of muscle-related biological processes being inhibited (Supplemental Fig.1B).

**Figure 2.**
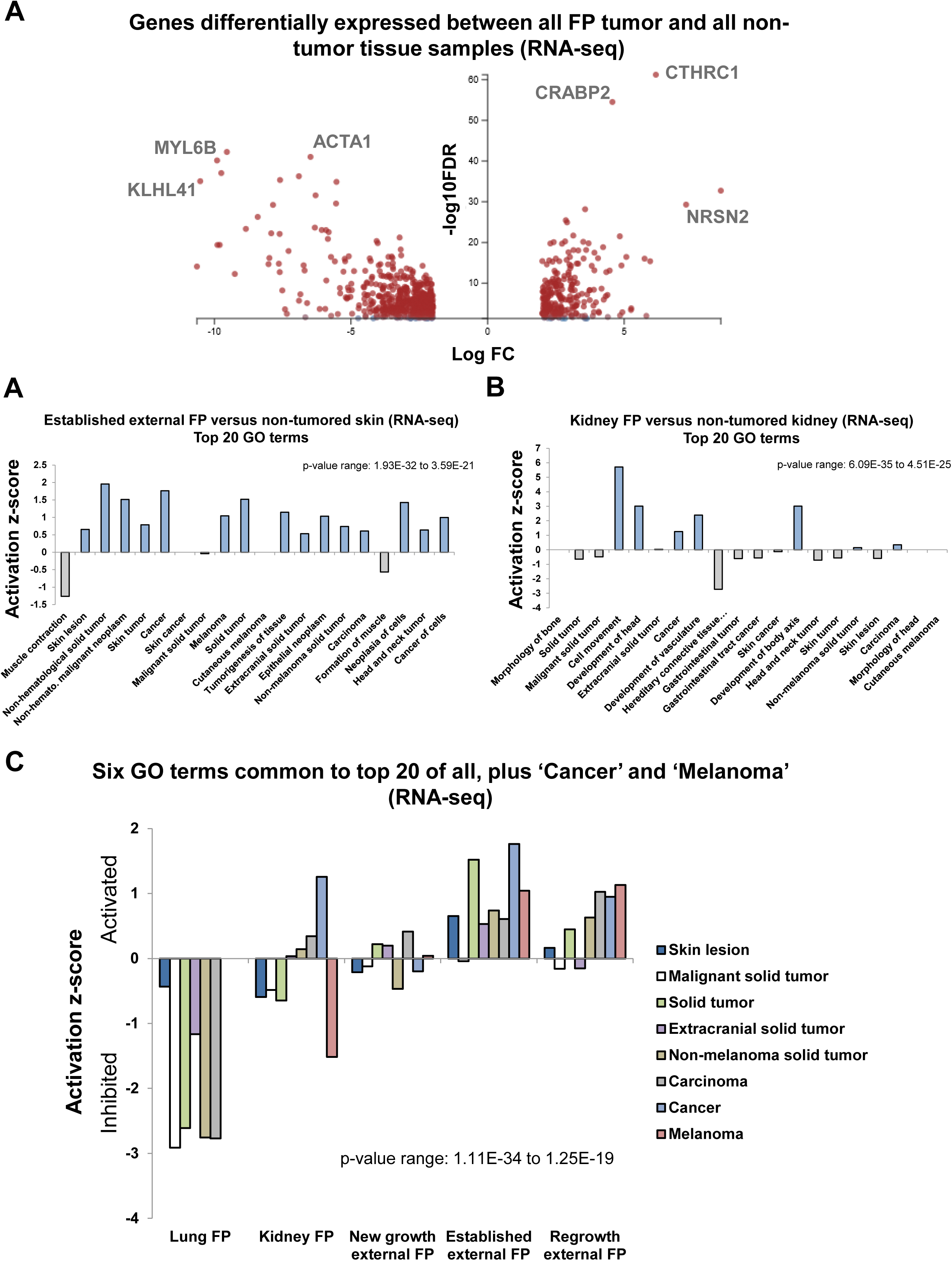
Gene Ontology (GO) term analyses of differentially expressed transcripts in each tumor type. **A)** Volcano plot of genes differentially expressed between fibropapillomatosis tumors (all types) and non-tumor tissue (all types) as determined by DESeq2 analysis of the RNA-seq samples. Transcripts were considered significant if passing the following cut-offs: adjusted p-value of < 0.05 and log_2_ fold change of >2 and <-2. Plot generated using Degust: interactive RNA-seq analysis (http://degust.erc.monash.edu/)^101^. **B, C)** Activation/inhibition z-scores of the top 20 disease-associated GO terms associated with transcripts differentially expressed in different types of fibropapillomatosis tumors (RNA-seq), as detected by IPA, ranked by p-value (calculated by right-tailed Fisher’s Exact Test, with Benjamini-Hochberg correction). **B)** Established external FP versus non-tumored skin. A total of 689 of the significant DEs were recognized by IPA and used in the analysis. **C)** Kidney FP versus non-tumored kidney tissue. A total of 618 of the significant DEs were recognized by IPA and used in the analysis. **D)** Activation/inhibition z-scores of the six GO terms common to the top 20 GO terms of all sample comparisons, plus the ‘Cancer’ and ‘Melanoma’ GO terms, as detected by IPA, ranked by p-value (calculated by right-tailed Fisher’s Exact Test, with Benjamini-Hochberg correction).

Twelve of the kidney fibropapillomatosis’s top 20 GO terms were cancer-associated, while all of the lung fibropapillomatosis’s GO terms were cancer-associated (Fig. 2C,D). Unusually, almost all of the lung fibropapillomatosis’s top 20 GO terms that had direction of activation/inhibition called were inhibited (Supplemental Fig.1C). This is in complete contrast to the external tumor types analyzed. A number of the kidney fibropapillomatosis’s cancer-associated GO terms were also inhibited (Fig. 2C). This is surprising, as the internal FP’s were not compared directly with external tumors, rather differential expression was called based on comparison with their patient matched non-tumor internal tissue samples (non-tumored lung and non-tumored kidney). To ensure no non-tumor sample related bias, we also directly called transcripts differentially expressed (DESeq2) between established external FP samples and internal lung FP or kidney FP samples (Supplemental Fig. 2A-C). This showed 11 cancer-associated GO terms (of the top 20 GO terms) to be activated in lung FP (Supplemental Fig. 2C). Only four of the top 20 GO terms were inhibited cancer-associated terms (Supplemental Fig. 2C). This suggests that the inhibition of cancer-associated GO terms seen when lung FP was compared with non-tumor lung (Supplemental Fig.1C) may be due to non-tumor tissue specific expression variation, rather than cancer-associated processes being inhibited in lung FP tumors. As demonstrated by the PCA plot, the expression profiles of the internal non-tumor tissue samples differed widely from the skin and external FP tumor samples (Fig. 1E). Overall, of the top 20 GO terms from each tumor compared with their matched non-tumor tissue type, six were common to every FP type sequenced, although their activation/inhibition status differed between internal and external tumors, as did their extent of activation across the three external tumor types (Fig. 2D).

To further determine what signaling events are prominent in driving early external tumor formation (new growth and post-surgical regrowth), we examined the top 200 (ranked by p-value) GO terms in early external tumors compared with established external tumors (Fig. 3A). Similar to the gene level analysis there was a very high degree of overlap between the three external tumor types. However, of the 200 GO terms, 22 were uniquely common to early new and regrowth tumors (Fig. 3A). Of these 22 GO terms, eight of them were associated with Leukocyte/lymphatic processes (Fig. 3B), suggesting a crucial role of immune response during initiation of FP tumor growth. Immune activation may be in response to the ChHV5 virus, or to the tumor cells themselves. All eight leukocyte/lymphatic process GO terms were strongly activated across the three external FP types (Fig. 3B), although for established external tumors these terms fell outside of the top 200 GO terms. Interestingly, two of these eight terms (cell movement of leukocytes and leukocyte migration) were called as statistically significant in internal tumors (although outside of the top 200. Fig. 3B). In kidney FP tumors both terms were strongly activated, mirroring the external tumors. Conversely however, in lung FP tumors both of these GO terms were inhibited (Fig. 3B), again highlighting differences between the molecular signaling events driving internal and external FP tumors. Interferon gamma, a cytokine critical to both innate and adaptive immunity, was also called as an inferred transcriptional regulator (ITR) in all five tumor types, being activated in every tumor type barring lung tumors where it was inhibited (Supplemental Fig. 2D)

**Figure 3.**
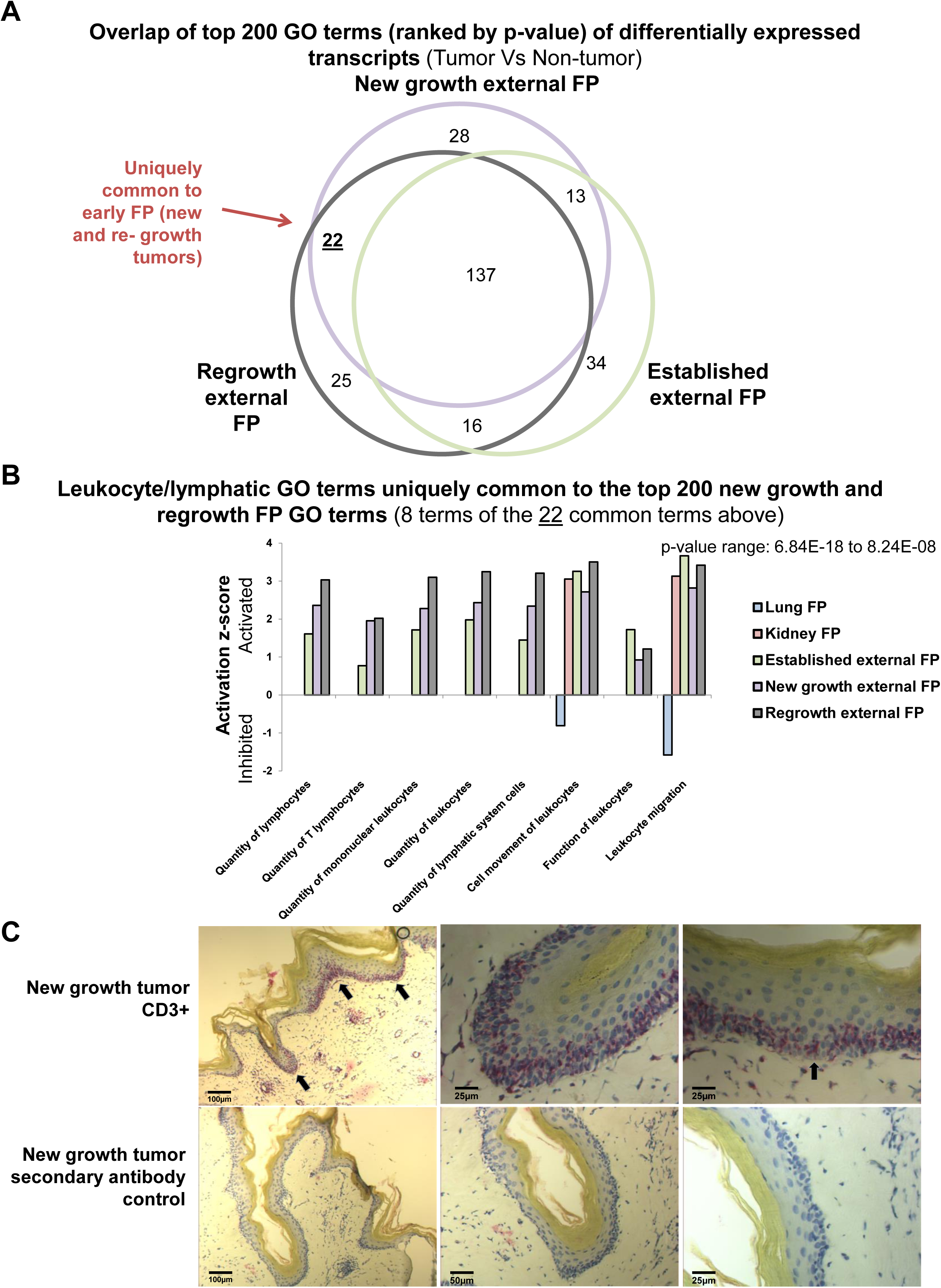
Transcriptomic- and histological-based immune profiling of fibropapillomatosis tumors. **A)** Overlap of the top 200 disease-associated GO terms associated with transcripts differentially expressed in different growth stages of external FP tumors (new growth, regrowth, established), as detected by IPA, ranked by p-value (calculated by right-tailed Fisher’s Exact Test, with Benjamini-Hochberg correction). Both activated and inhibited GO terms for each tumor type when it was compared to its healthy tissue source were included. Area-proportional Venn diagrams were generated using BioVenn (http://www.biovenn.nl/). **B)** Activation/inhibition z-scores for eight (the leukocyte/lymphatic-associated GO terms) out of the 22 disease GO terms uniquely common to the top 200 ranked GO terms of early fibropapillomatosis (new growth and regrowth, see Fig. 3A), shown for all tumor types. Some of these GO terms were called for lung, kidney and/or established external tumors (as shown), although they fell outside of the top 200 ranked GO terms called for these three tumor types. **C)** CD3 anti-body based staining of T lymphocyte infiltration in new growth tumor tissue, with alkaline-Phosphatase based secondary anti-body staining producing reddish/purple positive staining. For ease of visualization nuclei are counterstained with Hematoxylin (blue staining). Selected positive CD3 stained areas are indicated by black arrows.

Given the strong leukocyte/lymphocyte infiltration results from the tumor transcriptomics, we next assessed the host adaptive immune response by identifying CD3 positive T lymphocytes^56^ within fibropapillomatosis tumor tissue sections. This confirmed the transcriptomics findings, revealing that even in early stage FP tumors there is a high level of T lymphocyte infiltration (Fig. 3C). CD3 is an immunophenotypic cell marker, which is found only in T lymphocytes and is central to the formation of antigen-receptor interactions through the T cell receptor/CD3 complex ^56, 57^. CD3 positive staining was strongest in epidermal regions, where inclusion bodies (presumably due to lytic ChHV5) most commonly occur within FP tumors^12^. Together the transcriptomics and CD3 staining demonstrate that an immune response is mounted by the host (*C. mydas*), either to the tumor cells themselves, and/or more likely given the localization pattern to ChHV5 infection.

The disparate signaling events detected by the transcriptomics between external, lung and kidney tumors, potentially make it less likely that a single systemic anti-cancer therapeutic would prove effective against both external and internal tumors. However, to investigate whether any common therapeutically targetable oncogenic pathways exist between these tumor types we next compared their top 100 ITRs. IPA analysis infers the upstream transcriptional regulators responsible for the observed transcriptomic signatures by comparing the differential gene expression profiles to known regulator induced changes in its knowledgebase. Mirroring the gene level analysis, ITR analysis also showed very little overlap between the top 100 ITRs of established external, lung and kidney FP tumors (Fig. 4A). However, if a common therapeutically targetable vulnerability exists it should be located in the overlapping ITRs of these three FP tumor types. Therefore, we further investigated the 16 ITRs which all three FP types had in common (Fig. 4 A-C). These 16 ITRs represented nine genes and seven pharmaceutical compounds (drugs). Of the nine gene ITRs, almost all were activated across all five FP tumor types sequenced (external: established, new and regrowth, and internal: lung and kidney), with new growth external tumors tending to be the exception (Fig. 4C). These nine genes form a highly interconnected regulatory network (Fig. 4B), with 32 edges between the nodes and a protein-protein interaction enrichment p-value of 3.82e-07 (String). Interestingly, retinoic acid (RA) signaling was activated strongly in established external, external regrowth and kidney FP tumors (Fig. 4C). Retinoid therapy is widely used as an anti-cancer therapeutic and maintenance therapy for a number of human cancers (such as the pediatric cancer neuroblastoma^58^), although conversely RA signaling is known to be activated in some cancers. Retinoic acid is a widely available and inexpensive anti-cancer therapeutic, which would make it ideally suited for use in sea turtle rehabilitation facilities. Unfortunately the findings here suggest RA would be ineffective against FP. These transcriptomic findings explain the failure of early attempts to treat FP tumors and prevent regrowth using ectopic RA application (Supplemental Fig. 3A).

**Figure 4.**
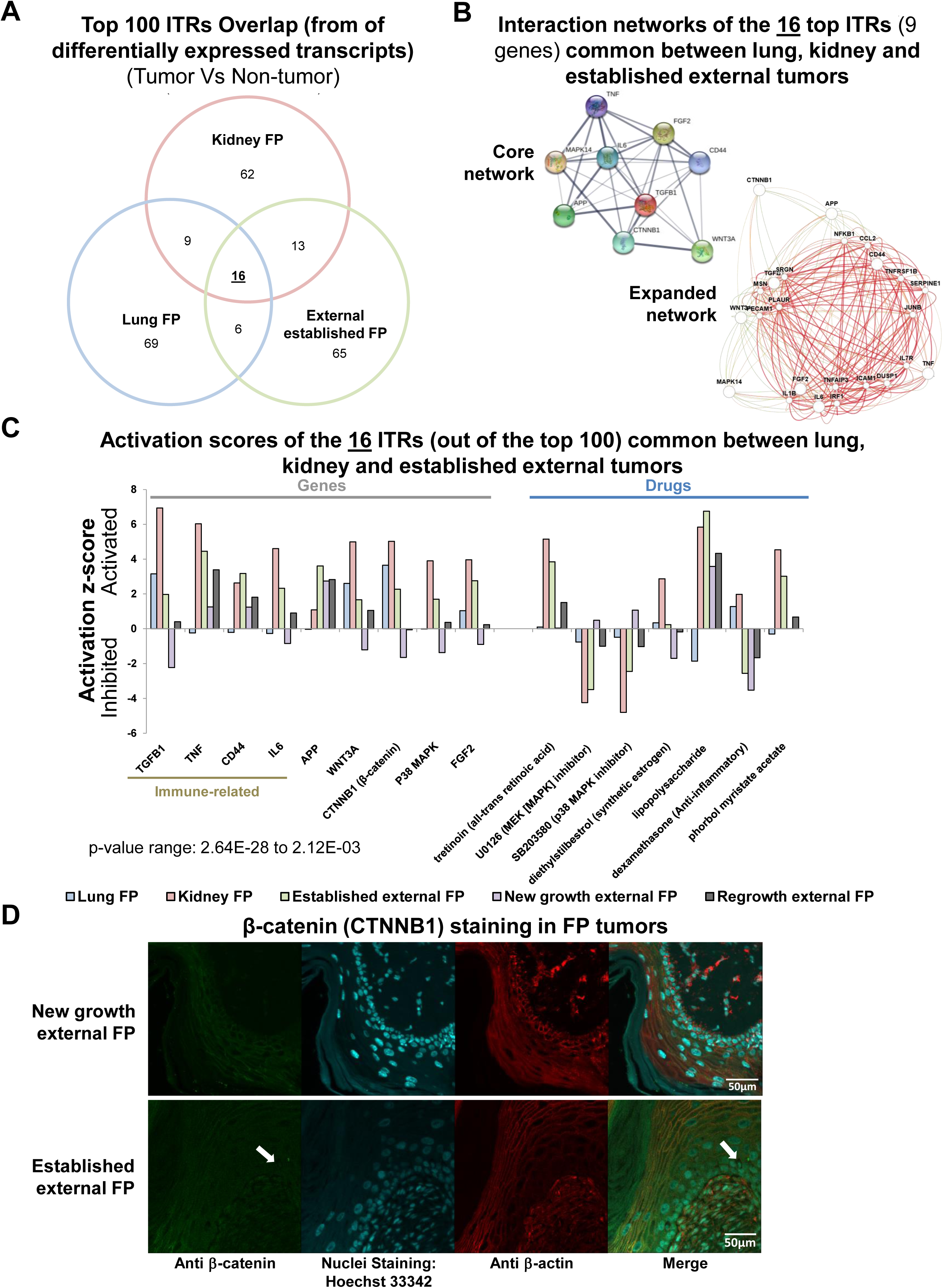
Transcriptional regulator analysis revealing the cellular signaling driving fibropapillomatosis and potential therapeutic targets. **A)** Overlap of the top 100 inferred transcriptional regulators (ITR) of the transcripts differentially expressed in different types of FP tumors (kidney FP, lung FP, external established FP) when compared to their respective non-tumored tissue sources, as detected by IPA, ranked by p-value (calculated by right-tailed Fisher’s Exact Test, with Benjamini-Hochberg correction). Area-proportional Venn diagrams were generated using BioVenn (http://www.biovenn.nl/). The 16 ITRs that were common to lung, kidney, and established external FP were selected for further analysis. **B)** Interaction networks of the top 16 ITRs (9 genes shown, 7 drugs excluded) common between lung, kidney and established external tumors. Core network generated by String^99^ (https://string-db.org/), expanded network generated by HumanBase (https://hb.flatironinstitute.org/). **C)** Activation/inhibition z-scores for the 16 ITRs common between lung, kidney, and established external tumors, shown for all tumor types. ITRs are segregated according to functional class, i.e. genes and drugs **D**) Anti-body based immunohistochemistry of new growth and established external tumor tissue. Tissue sections are stained with Anti β-catenin (an ITR called for 3 of the 5 tumor types, see Fig. 4C) and counter stained with Hoechst 33342 to visualize nuclei and Anti β-actin. Selected cells with nuclear (activated) β -catenin staining are indicated by white arrows.

The 16 ITRs (Fig. 4C) tended to fall into three main categories, canonical Wnt signaling, MAPK signaling and immune-related signaling (CD44, IL6, TNF, TGFβ). These pathways form part of an interlinked signaling network (Fig. 4B). Amyloid precursor protein (APP) may also be immune-related, as in addition to being implicated in neuron synapses and Alzheimer’s disease, it has also been implicated to be involved in antimicrobial activity and iron export. As predicted by the transcriptomics, β-catenin was located at the cellular membrane in new growth external FP tumors, indicating inactivation of Wnt/β-catenin signaling (Fig. 4D). Again in line with the ITR analysis, nuclear localization of β-catenin was present within other tumor types (Fig. 4D and Supplemental Fig. 4A), which is indicative of Wnt/β-catenin pathway activation. While nuclear β-catenin did occur in external established, regrowth, and internal tumors it was far from ubiquitous, suggesting intra-tumor heterogeneity in terms of Wnt signaling activation. Generally the transcriptomics revealed that each of the three main shared signaling pathways (Wnt signaling, MAPK signaling and immune-related signaling) were activated; inversely, but logically, the drug inhibitors of these pathways were inhibited (U0126, SB203580 and dexamethasone, Fig. 4C). This suggests that FP tumors (both external and internal) may be susceptible to treatment with inhibitors of these pathways.

### Transcriptionally-inferred molecular origins of fibropapillomatosis

To gain further insights into FP’s origins we employed larger scale network clustering analysis. Established tumors demonstrated clusters of ‘viral and inflammatory responses’, ‘inhibition of anoikis’ (programmed cell death of anchorage-dependent cells that detach from the extracellular matrix), ‘cellular senescence’ and ‘miRNA regulation’ (Fig. 5A). Interestingly, given FP’s environmental trigger(s), established external tumor’s highly interconnected ITR network (protein-protein interaction enrichment p-value <1.0E-16) also had signatures related to ‘cellular responses to organic substances and chemical stimulus’ (Supplemental Fig. 5A). Furthermore, pathways related to ‘Kaposi sarcoma-associated herpes virus infection’ (KEGG pathway analysis 1.20E-32) were also detected in established external FP (Supplemental Fig. 5A). Kaposi sarcoma is a human cancer which arises in immunocompromised patients, with lesions developing on skin, lymph nodes, or other organs, and is associated with a herpesvirus (human herpesvirus 8, HHV8). Like HHV8, the chelonid herpesvirus, ChHV5, is similarly postulated to drive FP tumor formation in sea turtles compromised by an external environmental or immunomodulatory trigger, a hypothesis which is strengthened by the viral and immune response signaling dynamics revealed by the transcriptomics. Immune-related processes featured in all five tumor types (Fig. 5A,B and Supplemental Fig. 6A-C). Although transcriptionally divergent to established external FP tumors, lung FP also showed an interconnected network (protein-protein interaction enrichment p-value <1.0E-16) with cellular immune response and organic substance and chemical response nodes (Supplemental Fig. 5B). Furthermore, clusters related to response to inorganic substances, metal ions, viruses and radiation were also detected (Fig. 5B). Indeed, ‘Quantity of Metal’ was called as a GO term and was activated for all tumor types, with the exception of kidney tumors in which it was not called (Supplemental Fig. 6D).

**Figure 5.**
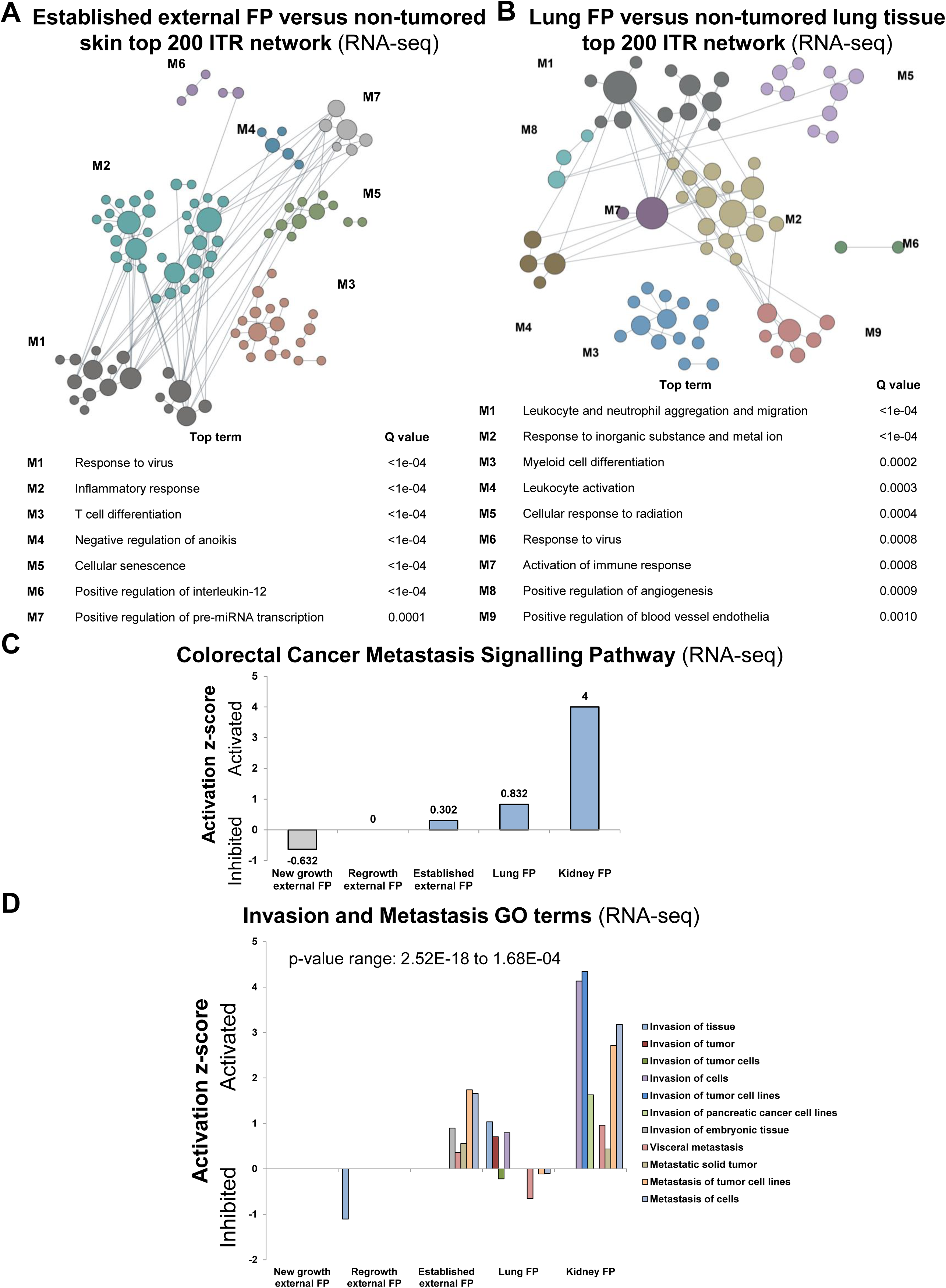
Network analysis of inferred transcriptional regulator analysis of fibropapillomatosis tumors. **A, B)** Network-based functional module discovery of the top 200 ranked ITRs (called by IPA) of **A)** established external and **B)** lung tumors generated by HumanBase (https://hb.flatironinstitute.org/). **C)** Activation/inhibition z-scores of the ‘Colorectal Cancer Metastasis Signaling’ gene ontology (GO) term associated with transcripts differentially expressed in different types of fibropapillomatosis tumors (kidney FP, lung FP, external FP) when compared to their respective non-tumored tissue sources, as detected by IPA. **D)** Activation/inhibition z-scores of 11 invasion and metastasis associated GO terms from transcripts differentially expressed in different types of fibropapillomatosis tumors (kidney FP, lung FP, external FP) when compared to their respective non-tumored tissue sources, as detected by IPA.

Pathway analysis (Fig. 5C) and GO term analysis (Fig. 5D) revealed a graded activation of metastatic-related signaling across the FP tumor types. FP tumors have been described as primary, despite the numerous tumors which regularly develop on each afflicted individual. However, to date no in-depth molecular analysis has been conducted to determine if all tumors on an individual are indeed primary, or if all or some of them (particularly internal visceral tumors) occur due to metastatic spread of a primary tumor. Our analysis suggests that the situation may not be so clear-cut, and therefore systematic phylogenetic/phylogenomic analysis of numerous tumors upon the same individual should be conducted. Our transcriptomics suggests that at a minimum, fibropapillomatosis tumors have a propensity to mutate towards the activation of metastatic pathways, with internal tumors showing stronger activation than external tumors (Fig. 5C,D). Early external FP tumors do not display metastatic signaling, rather such pathways are mildly inhibited (Fig. 5C,D). Established external FP show mild metastatic signaling activation, while internal tumors, particularly kidney tumors, show elevated activation of these pathways (Fig. 5C,D). It should be determined whether this activation is due to metastasis having occurred, or whether the propensity to metastatic activation falls short of complete metastasis.

### The genomic mutational landscape of disseminated FP tumors

Genome instability and mutation (host) is a hallmark of cancer ^59^, DNA replication and repair machinery malfunction frequently during oncogenesis leading to cancer cells having aberrant genomes. Such genome instability and mutations can range from single base pair changes to whole chromosome gains and deletions ^59^. Indeed, initial mutations within the genes controlling DNA replication and repair can be self-selecting leading to further cancer promoting genomic alterations. The mutational landscape of the host genomes of marine turtle fibropapillomatosis has yet to be explored, we therefore, next conducted whole genome sequencing (WGS, DNA-seq) of three fibropapillomatosis tumors and their patient-matched non-tumor tissue, to investigate whether they harbor genome instability in the form of copy number variations (CNVs), regions of the chromosome which have additional copies (gains) or which have been deleted (losses). Copy number variation analysis revealed that fibropapillomatosis tumors can harbor mutations, i.e. may not solely be driven by non-genomic events (e.g. viral, epigenetic or environmental). However, inter-tumor heterogeneity was observed, even between two tumors (one lung and one kidney) from the same individual (Figs 6A, Supplemental Fig. 7A). Copy number gains were present in the lung tumor (Fig. 6A), while no copy number variations were observed in the other two tumors (Supplemental Fig. 7A). The heterogeneous CNV landscape of the lung and kidney tumor from the same individual (patient 27-2017-Cm), suggests that these two tumors are not clonal (to each other) and did not arise as metastatic spread from the same original primary tumor. Although, it remains a possibility that the kidney tumor metastasized to the lung, only gaining CNVs after leaving the kidney.

**Figure 6.**
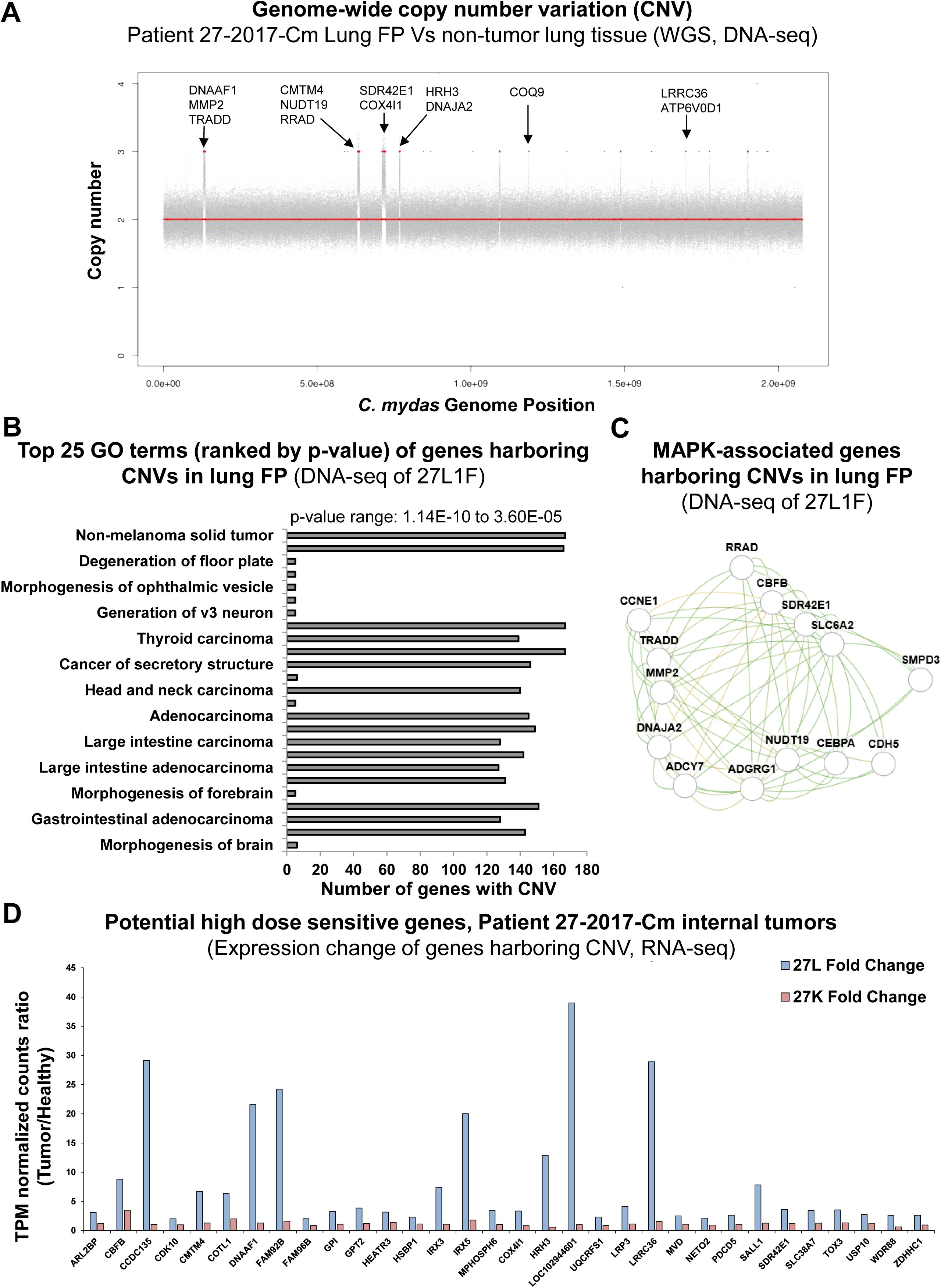
Genome-wide copy number variation (CNV) analysis of whole genome sequencing of patient-matched tumor and non-tumor tissue reveals possible oncogenic drivers in a lung tumor. **A)** Copy number variation analysis using the cn.MOPS package (Klambauer et al. 2012) implemented in R (https://www.r-project.org/). Tumored tissue was compared to non-tumored tissue from the same tissue source and same patient (27-2017-Cm). Each point shows a genomic segment 5000 bp long and its associated copy number within the tumor relative to its patient-matched non-tumor tissue, with a copy number of 2 being the null hypothesis for a diploid organism. Red data points indicate where the segmentation algorithm called a potential significant CNV. **B)** Top 25 disease-associated gene ontology (GO) terms, as detected by IPA, ranked by p-value (calculated by right-tailed Fisher-s Exact Test, with Benjamini-Hochberg correction) of the genes found within the potential significant CNV regions identified in 27-2017-Cm’s lung tumor. While 220 genes were extracted from the putative CNV regions, only 174 were mapped (human-centric) and included in the IPA analysis. **C)** Functional interaction network of the MAPK-associated genes harboring CNVs in 27-2017-Cm’s lung tumor. Network generated using HumanBase (https://hb.flatironinstitute.org/). **D)** Comparison of gene expression differences between kidney tumor (27K, copy number neutral) and lung tumor (27L, copy number gains) for potential highly dose sensitive genes found within the lung tumor CNV regions of 27-2017-Cm. Count ratios were generated by taking the transcripts per million (TPM) normalized read counts per gene from tumor samples and dividing that value by the normalized counts per gene from the patient-matched non-tumor tissue samples.

Next the CNVs occurring in Patient 27-2017-Cm’s lung tumor were mapped to the gene level revealing 228 genes (174 of which were recognized by IPA [human filter]) with copy number gains. When this CNV gene list was subjected to IPA analysis it was highly enriched for cancer-associated genes, with non-melanoma solid tumor (a human skin cancer) being the most enriched pathway (p-value 1.14E-10). This indicates that these CNVs may be involved in driving FP tumorigenesis and growth/progression (Fig. 6B), rather than solely being passenger mutations. Furthermore, 22 Wnt/β-catenin-related and 14 MAPK associated genes harbored copy number gains (Fig. 6C), with p38 MAPK signaling, ultraviolet radiation (UV)-induced MAPK signaling, MAP2K1, MAP3K4 and MAP3K8 called as pathways and ITRs of the CNV harboring gene list (IPA). That UV-induced MAPK signaling (particularly the SMPD3 gene) was called is particularly interesting as UV exposure is a putative environmental trigger of FP ^26, 30^. Additionally, there were epigenetic-related genes harboring copy number gains in the lung tumor, such as BRD7 which is involved in chromatin remodeling complexes. Genes harboring copy number gains in Patient 27-2017-Cm’s lung FP genome had corresponding expression gains when dosage analysis was performed using the corresponding RNA-seq data for this tumor (Fig. 6D, Supplemental Table 3), confirming that these mutational gains induced changes in gene expression. These mutated genes are also involved in the regulation of cell proliferation by regulating the mitotic cell cycle regulation and DNA replication checkpoint (Supplemental Figure 7B), dysregulation of these cycles is a key hallmark of cancer cells ^59^.

### ChHV5 is latent in FP tumors, including early stage and internal tumors

While further investigation of the genomic and environmental drivers of FP is warranted, the suspected causal relationship between ChHV5 and FP also requires further study. We demonstrated previously that ChHV5 was transcriptionally latent in established FP tumors ^26^. However, that study only examined a small number of established external tumors (seven) and did not investigate whether ChHV5 was lytic or latent in early stage, regrowth or internal tumors. We postulated that ChHV5 might be latent in established tumors, but more active during crucial early stage tumor initiation events, akin to the ‘hit and run’ hypothesis of viral oncogenesis ^26^. To investigate this hypothesis, we aligned the reads from the 90 RNA-seq samples above to the ChHV5 genome.

Across all FP samples, ChHV5 transcripts were low, and we detected no major switch to active (lytic) virus in either new growth FP or post-surgical regrowth FP (Fig. 7A). In fact, levels of ChHV5 transcripts were only marginally higher in FP tumors than they were in non-tumored tissue controls (Fig. 7A), though this was not significant for any of the FP tumor types compared with the non-tumor tissues. The only significant difference found in the level of viral RNA transcripts between any of the groups was between regrowth and established growth external tumors (Kruskal Wallis with Dunn-Bonferroni post hoc, p = 0.013). Furthermore, we detected no switch to active (lytic) viral transcription in internal tumors (Fig. 7A). Together this suggests that the role of lytic ChHV5 in FP is minimal and that if ChHV5 is contributing to FP oncogenesis, it is either occurring transiently, during very early tumorigenesis (before visible tumor appears), or through the expression of ChHV5 miRNAs (not assessed here), or that it is the latently expressed ChHV5 transcripts that are driving oncogenesis. We therefore next examined the individual gene level transcripts to determine which ChHV5 genes were transcriptionally active in FP tumors (albeit at relatively low levels). Samples were grouped into four types: non-tumor, external FP (including established, new growth and regrowth), kidney FP, and lung FP. Across all four sample types quite a consistent pattern of ChHV5 gene expression emerged (Fig. 7B, Supplemental Fig. 8A), with only 22 of ChHV5’s 104 genes showing levels of expression above 1 transcripts per kilobase million (TPM, Supplemental Table 2). Latency-associated genes such as F-LANA formed part of this 22 gene group, being consistently expressed across all tumor types (Figs 7B and Supplemental 8A). Given the paucity of lytic ChHV5, these 22 ChHV5 genes represent the most likely viral drivers able to contribute to FP initiation and ongoing tumor development and growth; therefore they warrant further functional investigation.

**Figure 7.**
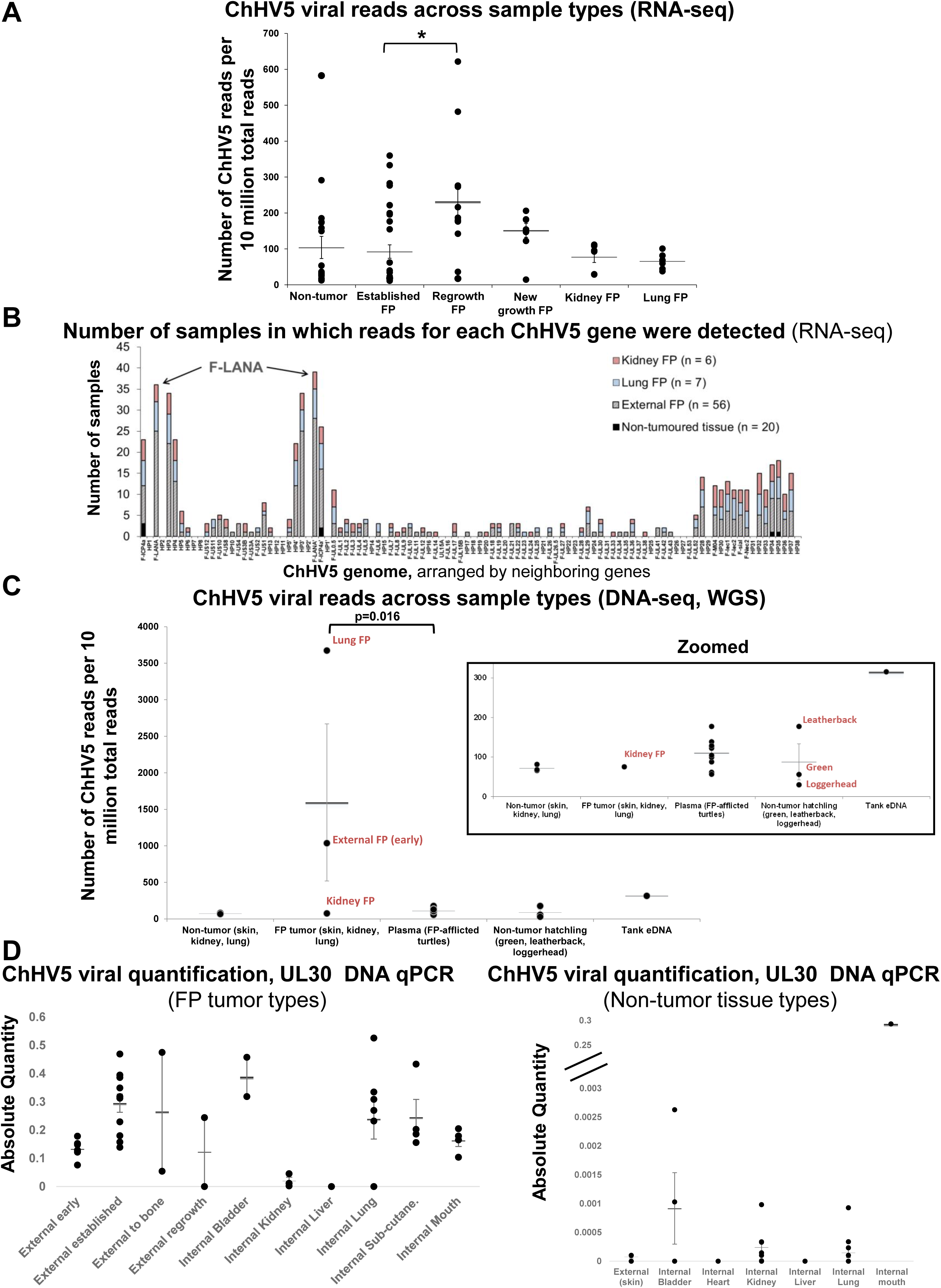
ChHV5 transcriptomics and genomics across tumor and non-tumor samples. **A)** ChHV5 expression across each sample type. Dot plot of the number of ChHV5 reads (RNA-seq) per 10 million total reads per sample. Line denotes mean, error bars denote standard error. Significant differences in averages between sample types were determined by a Kruskal Wallis with Dunn-Bonferroni post hoc and are denoted by an asterisk (*). Per sample type, non-tumor n = 20, established external tumor n = 35, regrowth external tumor n = 12, new growth external tumor n = 9, kidney tumor n = 6, lung tumor n = 7. **B)** Total number of samples in which reads (RNA-Seq) for each ChHV5 gene were detected. A gene was counted as detected if a sample had TPM-normalized counts > 0 for said ChHV5 gene. **C)** ChHV5 abundance (DNA-based) across each sample type. Dot plot of the number of ChHV5 reads (DNA-seq) per 10 million total reads across sample types. Line denotes mean, error bars denote standard error. Per sample type, non-tumor n = 3, fibropapillomatosis tumor n = 3, plasma from FP-afflicted turtles n = 10, tissue from non-tumor hatchlings n = 3, tank environmental DNA (eDNA) n = 1. **D)** ChHV5 viral quantification of a range of FP types (n = 43 samples) and non-tumor tissue (n = 36) samples from 13 individual patients using the UL30 DNA qPCR assay^25^ (Supplemental Table 4). Error bars denote standard deviation. Absolute quantity of ChHV5 was determined through a standard curve of known amounts (in picograms) of a UL30 gene fragment (Supplemental Table 5). Truncated x-axis label, internal sub-cutan. = internal sub-cutaneous.

Almost all samples with above 200 ChHV5 viral reads per 10 million total reads originated from just three patients (Supplemental Fig. 9A). Interestingly all three of these higher viral transcript patients were successfully rehabilitated and released. This prompted us to investigate whether there was any relationship between number of viral transcripts and rehabilitation outcome. Counter-intuitively, patients with positive outcome (release) on average had samples with statistically significantly higher ChHV5 transcripts (Mann-Whitney U Test, *p* = 0.03), while those patients that died in care or were euthanized due to advanced disease actually had lower ChHV5 transcripts (Supplemental Fig. 9B). Even when internal tissue samples were removed from the analysis (as all internal samples originated only from the deceased/euthanized category), there remained a significant difference between the two groups (Mann-Whitney U Test, *p* = 0.0143).

### High ChHV5 viral loads, predominantly arise from latent virus

Given the low level of ChHV5 transcripts in FP tumors, we next used whole genome sequencing (WGS/DNA-seq) to determine the viral load of ChHV5 to determine whether the low number of transcripts arises due to a lack of virus, or whether large quantities of virus are present within FP, but that the majority of these are latent (not undergoing active viral replication and transcription). By not conducting any viral enrichment steps, the resulting read numbers give a more reliable indication of the relative abundance of viral DNA compared with host DNA (all within a single sample/library). Viral DNA sequencing reads in the FP tumors covered a broad range. Interestingly, the quantity of viral DNA reads in FP tumors was not significantly different when compared to any other group, with the exception of the plasma samples (Fig. 7C). The lung and kidney tumor DNA-seq samples (also used for host genome CNV analysis) from the same Patient 27-2017-Cm had dramatically different viral loads (Table 1). The early growth FP tumor from ‘Lilac’ (Patient 25-2018-Cm) had an intermediate viral load 1,036 ChHV5 per 10 million total reads (RPTMT, Table 1 and Fig. 7C). Despite the high level of ChHV5 DNA sequencing reads in the lung tumor (3,673 RPTMT), this same tumor only had a very low level of ChHV5 RNA sequencing reads (68 RPTMT), more closely resembling the read numbers of the non-tumor tissues (Table 1). This suggests that rapid proliferation of host tumor cells, infected with latent virus is the primary driver of the high viral loads sometimes observed in FP tumors (some FP tumors can double in size in less than two weeks^46^).

**Table 1.** Comparison of viral sequencing reads per tumor and non-tumor tissue at the DNA and RNA level. Number of reads per 10 million total reads (RPTMT) are shown for each sample.

| Sample | DNA viral load (RPTMT) | RNA viral reads (RPTMT) |
| --- | --- | --- |
| <u>Patient 27-2017-Cm:</u> |  |  |
| Lung FP tumor | 3,673 | 68 |
| Lung non-tumor | 68 | 36 |
| Kidney FP tumor | 75 | 30 |
| Kidney non-tumor | 66 | 26 |
| <u>Patient 25-2018-Cm ('Lilac'):</u> |  |  |
| External new growth FP tumor | 1,036 | - |
| Skin non-tumor | 81 | - |

The 27-2017-Cm kidney tumor had minimal host CNVs and minimal viral load (Table 1 and Fig 7C) suggesting that the tumor must be driven by other oncogenic mechanisms such as point mutations, epigenetic changes, or transcriptional/translational aberrations. The viral load of this kidney tumor was within the range of non-tumored tissue, plasma from FP-afflicted turtles and tissue from non-tumored hatchlings (Table 1 and Fig. 7C).

When DNA-based viral reads (Fig. 7C) are compared with viral reads from the RNA transcripts (Fig. 7A) of the 90 RNA-seq samples, the highest viral transcript load observed was approximately 600 RPTMT (one FP and one non-tumor sample), while the highest viral DNA load observed was 3,673 RPTMT. Although a high viral DNA load did not equate to high viral transcription (RNA) in the same sample (Table 1). As expected, WGS reads were dispersed across the entire ChHV5 genome (Supplemental Fig. 10A), not being restricted to ChHV5’s transcriptionally active regions (Supplemental Fig. 8A). This confirms that the limited transcriptional signature is genuine and not due to a sequencing artifact. Together the whole ChHV5 genome-level and gene-level TPM analysis highlights the marked difference in reads between viral DNA presence and viral RNA transcription in FP tumors. Conversely, the range of viral DNA and viral RNA within non-tumored tissue was largely overlapping suggesting that the ChHV5 present in non-tumored samples may be more likely to be transcriptionally active (Table 1 and Figs 7A, 7C).

We next examined the ChHV5 viral loads (viral DNA-based qPCR, ChHV5 UL 30 assay) in a wider cohort of internal tumor and matched non-tumor types. Lung and bladder FP tumors consistently had the highest viral loads (Fig. 7D), while of the non-tumor tissue types assayed, bladder tissue also showed some of the highest viral loads, with detected levels being higher than in some of the FP tumors (Fig. 7D). These results have implications for the spread of ChHV5, as urine is a potential source of ChHV5 transmission ^32, 33^.

### ChHV5 may be vertically transmitted from mother to offspring

Interestingly, the range of ChHV5 detected in hatchlings (one *C. mydas*, one loggerhead and one leatherback) overlapped the range of ChHV5 in FP-afflicted non-tumor tissue, the FP kidney tumor and FP-afflicted blood plasma samples (Fig. 7C). Of the three species, the leatherback sample had the highest number of ChHV5 reads. If hatchlings are already infected with ChHV5 (vertical transmission), this has serious implications for any potential population-level vaccination based mitigation strategies. On beach (immediate nest emergence^60^) sampling and sequencing-based ChHV5 detection should be conducted to confirm this finding and to determine the prevalence of ChHV5 infected hatchlings.

### Detection of ChHV5 shedding into the water column by environmental DNA (eDNA) approaches, reveals novel shedding dynamics and tumor burden correlations

A number of potential routes for ChHV5 transmission have been postulated, including via vectors such as *Ozobranchus* leeches^34^, which are commonly found on fibropapillomatosis-afflicted turtles, often at high density within the crevices of external fibropapillomatosis tumors (Fig. 8A). In line with previous studies^34^, we confirmed that ChHV5 could specifically be detected from DNA extracted from leeches which had fed on FP-afflicted turtles (Fig. 8A). Another potential mode of transmission is direct ChHV5 shedding into the environment. Direct viral shedding as a route of transmission is thus far only supported by indirect evidence, i.e. cloacal swabs, urine, ocular, oral and nasal secretions^32, 33, 61^, viral inclusion bodies near the surface of external tumors ^31^ and the elevated levels of ChHV5 detected in bladder tissue (Fig. 7D). Direct detection of ChHV5 in the water column is lacking. Therefore, we employed environmental DNA (eDNA)-based approaches coupled with qPCR and next-generation sequencing to detect ChHV5 in patient tank water. Such novel approaches can aid in answering previously intractable questions about direct ChHV5 transmission such as presence, abundance, and persistence in the marine environment. First, we determined that the presence of ChHV5 could be readily detected in eDNA extracted from tank sea water, using either an established^25^ UL30 qPCR assay or next-generation sequencing (Figs 7C, 8B-E). Furthermore, not only was ChHV5 detectible, it was quantifiable allowing us to make comparisons between individual tanks and across time (Fig. 8A-D). The level of detectible virus in patient tank water was positively correlated (R² = 0.5431, p = 0.0002, df = 19) to the tumor burden (as assessed by total tumor surface area) of the patient(s) housed in that tank (Fig. 8A). Patients with large well-established tumors shed more virus into tank water than those with small new-growth tumors. As FP tumors were surgically removed, so did the level of ChHV5 in patient tanks drop (Fig. 8C,D), suggesting that the tumors are the primary source of environmental ChHV5 (either through direct tumor shedding, or migration of virus throughout the body and excretion in bodily fluids). Additionally, the quantity of ChHV5 eDNA in tank water was positively correlated with the *C. mydas* (green turtle) eDNA level (R² = 0.66, p = 0.00001, df = 19), as detected by a custom *C. mydas* 16srRNA DNA assay (Fig. 8E).

**Figure 8.**
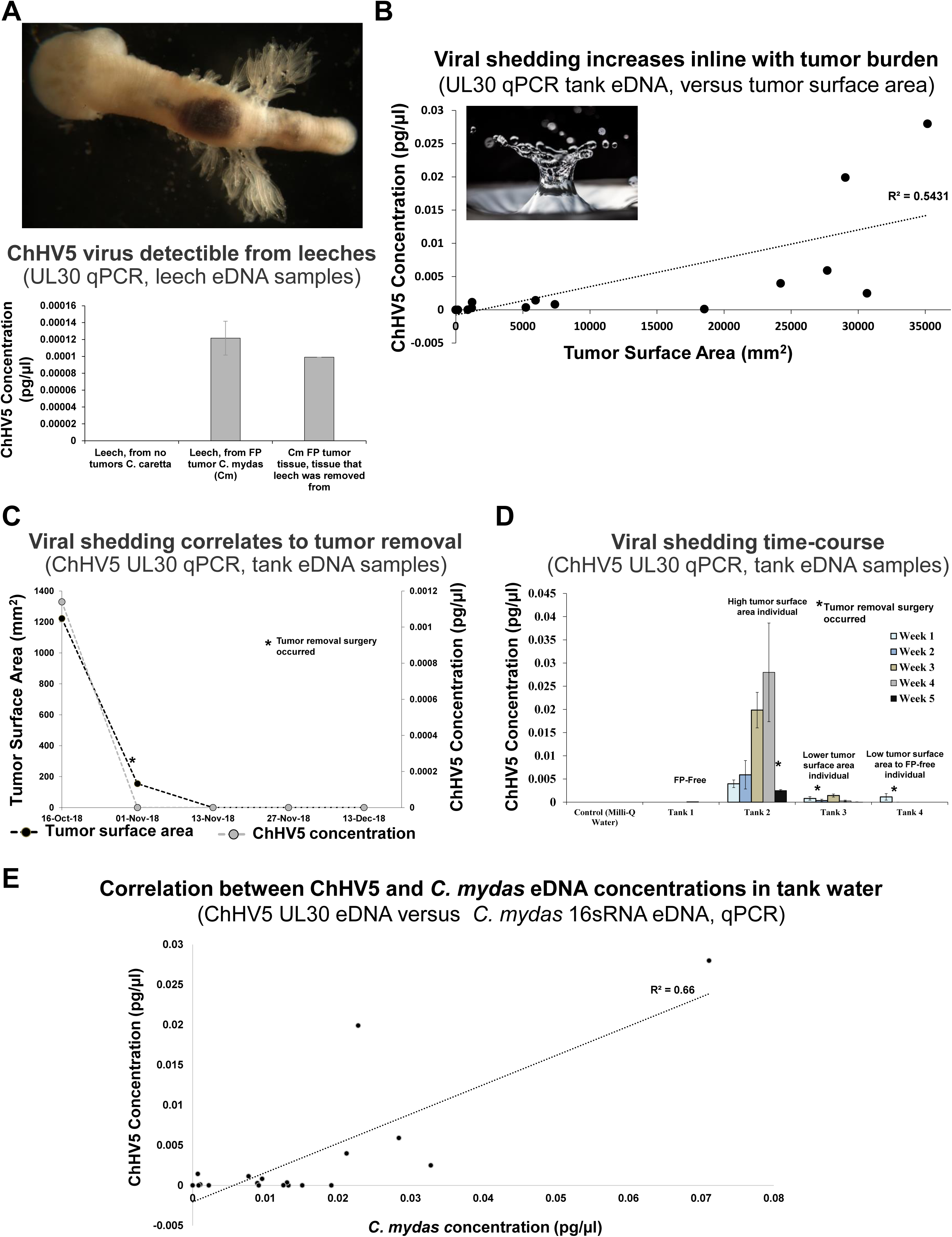
Environmental DNA (eDNA)-based detection, quantification and monitoring of ChHV5 viral shedding into patient tank water. **A)** Top, image of a marine leech removed from surface of a fibropapillomatosis tumor, with dark red *C. mydas* blood pellet visible. Bottom, detection and quantification of ChHV5 UL30 gene DNA by qPCR, using leeches as proxy eDNA samples (whole leech lysis and DNA extraction). Error bars denote standard deviation. **B) C**orrelation of individual patient tumor surface area (mm^2^) and the concentration of ChHV5 virus being shed into their tank water, as detected by qPCR (positive correlation, R² = 0.5431, p = 0.0002, df = 19). Background water drop image licensed under CC BY 2.0**C** (“Water Sculptures” by Tom Bullock, https://ccsearch.creativecommons.org/photos/6ccaf9a6-6a31-4aff-82fc-a071066fb689). **C)** Time-course of tumor surface area changes (surgical removal) and concentration of ChHV5 shed into tank water, as detected by qPCR. **D)** Time-course of ChHV5 viral shedding into four individual patient tanks, as detected by qPCR. Tumor removal surgery events are denoted by an asterisk. Note: the ChHV5 eDNA detected in tank 1 in week 4 was due to a second patient (FP-positive) being added to that tank for a single week, due to the rehabilitation needs of the hospital. Error bars denote standard deviation. **E)** Correlation of *Chelonia mydas* eDNA shedding and ChHV5 eDNA shedding (positive correlation, R² = 0.66, p = 0.00001, df = 19), both *C. mydas* (16srRNA gene assay) and ChHV5 (UL30 gene assay) eDNA were detected by qPCR.

## Discussion

### Transcriptional and mutational origins and therapeutic vulnerabilities of fibropapillomatosis

Wildlife pathogens have been shown to exacerbate the effects of environmental degradation, habitat loss and the climate emergency on population levels, potentially leading to local and global extinctions^2, 62–65^. Furthermore, as the risk of extinction increases for a given species the detrimental effect of disease on the population worsens^66^. Anthropogenic activities are not only stressing habitats, but the rapid environmental changes induced by these activities are likely increasing cancer rates in wild populations^67^. Human-induced perturbations of inshore marine environments have also been implicated as a co-trigger of the fibropapillomatosis tumor epizootic in green sea turtles^7, 29^. It is thought that environmental changes are key to conferring oncogenicity upon ChHV5, potentially through compromising or modulating the turtle’s ability to respond to the viral infection. We have demonstrated activation of immune-related signaling in FP tumors and shown localized CD3+ T-cell infiltration within new growth tumors. That the host still mounts an immune response within tumors is interesting given the previous links between FP-afflicted turtles and immunosuppression ^7, 11, 68, 69^. However, varying studies have found contradictory immunosuppression findings between FP and non-FP-afflicted individuals, for example no differences were observed in leukocyte activity between *C. mydas* turtles with and without FP, when blood samples were tested via flow cytometry^7, 10, 11, 70, 71^. Furthermore, not all turtles with FP are immunosuppressed and it is currently not clear whether immunosuppression in FP-afflicted turtles, when it does occur, is a cause or consequence of the disease ^7, 10, 11, 71^. Our transcriptomics and immunohistochemistry show that lymphocytes do infiltrate and mount an immune response within FP tumors. It will be important to elucidate why such a response is not sufficient at preventing initial tumorigenesis or preventing continued tumor growth. It should also be determined in the cases where spontaneous tumor regression does occasionally occur in non-heavily afflicted FP individuals whether regression is a consequence of elevated lymphocyte infiltration and activation^7, 10, 12, 72^. Further elucidation of the immune status of FP-afflicted individuals and the temporal dynamics and extent of immune responses will be crucial to determining whether the environmental co-trigger(s) of FP promote tumorigenesis through immunosuppression, be it systemic or localized.

Currently, no treatments exist for internal tumors with such turtles having to be euthanized regardless of their health status otherwise. Furthermore, surgical excision of FP tumors often results in high rates of tumor regrowth/recurrence^11, 26, 46^. Therefore, it is imperative that chemotherapeutic approaches are developed to augment surgical removal of external tumors and to provide first-line therapy for applicable internal tumors. We show that the molecular drivers of external and internal FP tumors are largely transcriptionally distinct. However, transcriptomic profiling and whole genome sequencing (CNV analysis) of FP tumors revealed that MAPK, Wnt and TGFβ pathway inhibitors are putative therapies for both external and internal FP, which could be targeted to simultaneously reduce tumor burdens (e.g. systemic targeting via oral dosing).

Although kinase inhibitors have been used extensively in human cancers, their application in veterinary medicine in mostly restricted to dogs and cats and only incidentally in other animals^73^. While Wnt signaling is heavily implicated in a wide range of human cancers, no clinically approved therapeutic yet exists, although clinical trials are ongoing^74, 75^. Similarly, TGFβ targeting therapeutics are undergoing clinical trials^76^. The MAPK/ERK pathway is involved in cell proliferation and survival and is implicated in carcinogenesis^77^. As such, a number of MAPK pathway drugs have been developed, such as MEK and BRAF inhibitors which are most commonly used to treat melanoma. MAPK/ERK inhibitors have greatly improved the life expectancy of patients with malignant melanoma, but acquired resistance almost inevitably occurs. Additionally, kinase inhibitors may cause serious adverse events for therapy recipients, with up to 46% of patients reported with grade 3-4 serious adverse events^78^. Worrying aspect of adverse effects with BRAF/MEK inhibitors is the occurrence of skin toxicities that also include secondary malignancies, such as keratoacanthoma and cutaneous squamous cell carcinoma^79^. RAF inhibitors may also cooperate with human papilloma virus to promote the initiation of cutaneous tumors^80, 81^. Given the complexities of patient responses to MAPK targeting therapeutics, other common pathways between internal and external tumors could be prioritized, or therapeutics selected to separately target external and internal FP tumors.

The novel findings of this study offer somewhat conflicting evidence about the nature of the relationship between internal and external tumors (i.e. whether they arise from primary or metastatic processes). Whole transcriptome profiling indicated that internal tumors are more closely-related to the tissue type in which they are found (presumptive tissue of origin), suggesting that they arise from the *de novo* transformation of these tissues and not as disseminated metastases from external tumors. However, skin cancer-related pathways/GO terms were called for internal tumors both from the differentially expressed genes (RNA-seq) and from the genes harboring copy number variation at the DNA level. Additionally, there was a trend towards increasing activation of metastasis-associated signaling particularly for kidney tumors. If ChHV5 is responsible for inducing *de novo* tumor formation in internal organs, then the limited expression and load of ChHV5 in some internal tumors is highly irregular. Similarly, the paucity of elevated levels of viral transcripts across all tumor samples suggests that lytic virus may not be driving the establishment of numerous primary tumors within the same individual. If ChHV5 truly is essential to the initiation of FP tumors and not just an opportunistic pathogen, it may be acting only through latently expressed genes (or microRNAs), or there may be an extremely limited temporal window during tumor initiation (tumorigenesis) in which lytic virus is more abundant (i.e. the viral hit and run hypothesis^82^), something we found no evidence for in any of the early stage tumors that were sequenced.

It is possible that a more complex situation exists with some internal tumors arising *as de novo* tumors while others within the same individual are due to metastasis. Given the presence of mutations within FP tumor genomes and their heterogeneity revealed here, in depth phylogenomics (clone history reconstruction) should be highly feasible, and its application could establish the clonal relationship between multiple tumors within the same individuals, including internal tumors. This would definitively address whether any FP tumor types arise as a consequence of metastatic spread. Given the heightened propensity of kidney tumors towards metastatic signaling pathway activation, an intriguing hypothesis would be that lung tissue receives sufficient air contact to enable lytic viral activation and *de novo* tumorigenesis, whereas kidney tissue does not. It was shown that in laboratory conditions ChHV5 would only undergo lytic reproduction within infected host cells when in direct contact with air ^48^. Supporting this, our whole genome sequencing and qPCR revealed that lung tumors tended to have much higher viral loads (DNA-based assessment) than kidney tumors (Fig. 7C,D), although there was no significant difference between the number of viral transcripts (RNA-seq) from lung and kidney tumors (Fig. 7A). Expanding similar genomics and transcriptomics approaches to tumors of other internal organ types would help elucidate whether there is a relationship between the level of the tissue’s air exposure and viral load or lytic virus level.

### The role of ChHV5 in tumor progression

The propensity of ChHV5 to only reproduce lytically (under laboratory settings) when infected host cells were in direct contact with air^31^, may also account for the localization of inclusion bodies within the epidermis of FP tumors^12^. It is possible that this feature could be an advantageous shedding strategy, as air contacting tissues are predominantly the skin surface and lungs, where shed virus would have a higher likelihood of successfully dispersing to other individuals. In line with this, we have shown that the level of ChHV5 in the water column positively correlates with level of external tumor surface area of patients.

Interestingly, when assessed in relation to patient survival and release, the presence of higher levels of ChHV5 RNA transcripts was actually prognostic for successful outcomes (Supplemental Fig. 9B). Patients having fewer viral transcripts were more likely to die or be euthanized during the course of their rehabilitation. Currently, due to the lack of treatment options (surgery is hampered by *C. mydas’s* hard plastron and carapace) any turtles found to be afflicted with internal tumors in Florida’s rehabilitation facilities are normally humanely euthanized^18^. Persistent aggressive post-surgical regrowth can also lead to the decision to euthanize a FP-afflicted patient. If ChHV5 is acting as an opportunistic pathogen in FP one might expect higher viral transcription in poor outcome patients, yet if ChHV5 is a driver of FP tumors it would also be expected that poor outcome patients would have higher levels of viral transcription (either lytic or latent associated gene transcription). Neither of these was the case for the patients whose tumors we sequenced. It would be informative to establish whether a similar relationship exists for viral load (DNA-based assessment) and outcome, or whether it is unique to viral transcription.

We reveal here that tumor-specific genome level mutations do occur within FP tumors, demonstrating that inter-tumor heterogeneity exists between two internal tumors from the same patient. Tumors harbored copy number gains in a variety of gene types and these were strongly enriched for cancer-associated functions. The mutational landscape of fibropapillomatosis, has important implications for understanding the disease’s molecular drivers and identifying potential therapeutic targets^2, 54^, and the identification of its environmental drivers (particularly single nucleotide polymorphism mutations)^83, 84^. Furthermore, better understanding of FP’s mutational landscape can help elucidate inter- and intra-tumor heterogeneity, the relationship between tumors within the same individual, and reveal the human cancer types which FP’s mutational spectra most resembles, informing the disease’s etiology and future treatment strategies^30^. Expanded whole genome-based mutational analysis may help explain how host cells can evolve and or rewire to become tumorigenic in the absence of high lytic virus loads, observed here.

Our findings demonstrate that latent ChHV5 predominantly accounts for the viral loads detected in FP tumors. Our genomic results show that the majority of ChHV5 replication in external and internal FP tumors is most likely driven through passive latent viral replication mechanisms, with the viral genomes being copied as the host tumor cells divide. Fibropapillomatosis tumors have been shown to grow rapidly ^46^. Our genomic analysis is consistent with the relative lack of lytic virus found within FP tumors by histological and single gene approaches^31, 34^.

### Mechanisms of viral exposure

ChHV5 DNA was identified in *C. mydas*, loggerhead (*Carretta carretta*), and leatherback (*Dermochelys coriacea*) hatchlings, and their abundance overlapped the range (no significant difference) seen in FP-afflicted non-tumor tissue, an FP kidney tumor and FP-afflicted blood plasma samples. Although it should be noted that ChHV5 genome coverage was not as complete in these hatchling samples as for tumor samples (data not shown), a key objective should be to confirm whether vertical transmission of the virus occurs from mother to offspring. This finding conflicts with the predominant hypothesis that ChHV5 infection is acquired only after juvenile turtles recruit to nearshore habitats^53, 85^. The occurrence of *in ovo* vertical transmission would have serious implications for the consideration of any potential future population-level vaccination-based mitigation strategies. On beach (immediate nest emergence^60^), or pre-hatching sampling and sequencing-based ChHV5 detection should be conducted to confirm this finding and to determine the prevalence of ChHV5 infected hatchlings.

Accurate detection and monitoring of wildlife pathogens with the capacity to impact species survival is essential to devise and implement appropriate mitigation policies^45^. Pathogenic viruses have the ability to infect susceptible species without presenting obvious symptoms^45^. Consequently, accurate, sensitive, non-invasive techniques are required to detect and monitor pathogens and pathogenic vectors outside of host systems^45, 86^. eDNA approaches have been shown to detect aquatic pathogens earlier than traditional methods and provide advanced warning of infection and mass mortality events^40, 45^. Recently, *Ranaviruses*, have been detected by eDNA analysis in environmental water samples from susceptible host species habitats, and demonstrated a strong relationship between environmentally-shed viral load and host tissue viral load^45^. Our application of eDNA approaches to detect ChHV5 in sea water is particularly significant given that unlike more immediate acting pathogens, there is generally a long lag time (years or decades) between infection and tumor formation^87–89^ for pathogen-induced cancers. This delay can hamper our understanding of infection dynamics and initiating events, making the ability for ongoing monitoring and surveillance critical.

ChHV5 can infect endangered sea turtle species without presenting signs of infection until environmental co-factors trigger transformation and oncogenesis; therefore diagnostic techniques capable of detecting the pathogen outside hosts is vital^33^. It has previously been asserted that only a small percentage of tumors shed virus (7% of tumors in 35% of individuals)^31^. However, our eDNA-based monitoring of viral shedding demonstrated that ChHV5 could be detected even in the tanks of turtles with low tumor burdens and that ChHV5 shedding positively correlated to tumor burden. This suggests that previous attempts to estimate shedding using localized approaches such as number of epidermal intranuclear inclusion bodies in tumor sections may underestimate the level of viral shedding occurring, with in-water ChHV5 quantification having improved detection limits than previous approaches. The future use of sensitive, novel molecular technologies such as eDNA may allow for the early detection of ChHV5 presence in the environment of vulnerable populations, and enable further research into the etiology, host species transmission and disease ecology of FP.

Our ChHV5 eDNA findings likely represent the first instance of time-course environmental DNA quantification of marine pathogen release from individual patients. This unique tool to dynamically track environmental viral shedding in response to disease progression and clinical interventions such as surgery and drug treatment will greatly enhance our ability to address fundamental questions relating to this disease, as well as to design evidence-based rational management strategies (e.g. containment/isolation policies). Such approaches can also improve our understanding of how environmental and clinical factors influence viral release and spread. The ability to perform eDNA-based quantitative viral shedding monitoring will also further enhance the utility of fibropapillomatosis as a model not just to better understand this wildlife epizootic, but also to address unresolved questions relating to viral shedding for other animal and human pathogen-induced cancers.

### Summary

Taken together, our results provide genome-level evidence for the complex relationship between external, new growth, established, post-surgical regrowth, and internal visceral tumors. They reveal the paucity of lytic ChHV5 replication within tumors, even those harboring high viral loads, as well as the heterogeneous mutational landscape of FP genomes and the host immune responses mounted within FP tumors. The application of precision oncology, genomic approaches, and novel technologies such as eDNA-based pathogen monitoring can assist in determining the molecular events underpinning FP tumor development, viral shedding and transmission dynamics, and enable the rational design of novel therapeutic interventions (such as pharmacological disruption of MAPK and TGFβ oncogenic signaling) and clinical management strategies. Importantly, the adoption of such approaches can elucidate the specific triggers of FP and the precise mechanisms through which these viral and environmental triggers are driving the fibropapillomatosis disease epizootic in marine turtles.

## Materials and Methods

### Tissue sampling

Sampling was carried out under permit number MTP-20–236 from the Florida Fish and Wildlife Conservation Commission and with the ethical approval from the University of Florida’s Institutional Animal Care and Use Committee (IACUC). External fibropapillomatosis (FP) tumors were surgically removed and punch biopsies taken of non-tumored areas as previously described ^26^. Internal tissue samples (tumors and non-tumor tissue) were obtained from animals during necropsies conducted immediately after euthanasia. Note that no animal was euthanized for the purposes of this study, but rehabilitating sea turtles determined to harbor internal tumors are currently euthanized in Florida as no treatment yet exists for internal tumors, and additional complications arising from surgery and other health concerns sometimes necessitates the humane euthanization of marine turtles in rehabilitation. Internal tissue samples were treated the same as the external samples. All samples were obtained from juvenile *C. mydas*, as this stage is the most commonly afflicted by the disease. Sex is not readily determinable in juveniles, but was provided for individuals that were euthanized due to internal tumors or other complications and in which necropsies were performed, or for individuals that were endoscoped (KARL STORZ, Multi-Purpose Rigid Endoscope for small animals) as part of their rehabilitative care (see Supplemental Table 1). Samples were stored until extraction in RNA-later (Qiagen) at - 80°C, according to manufacturer’s instructions.

### RNA and DNA Isolation, library preparation and sequencing from tissue samples

For RNA-Seq samples, total RNA was extracted using either an RNeasy Fibrous Tissue kit (Qiagen, Cat No. 74704) or RNeasy Plus kit (Qiagen, Cat No. 74134) with column-based genomic DNA removal, according to manufacturer’s instructions. Ninety RNA samples, comprising 70 fibropapillomatosis tumor samples and 20 non-tumor samples from 12 juvenile green turtles which had stranded in Northern Florida, were used for sequencing. Samples were further categorized by tissue type, as well as growth profile for the external tumors only (see Supplemental Table 1). Sequencing libraries were generated from 500 ng of total RNA using the NEBNext Ultra RNA Library Prep Kit for Illumina (New England Biolabs, Cat No. E7530), including polyA selection, according to manufacturer’s protocol. Size and purity of the libraries were analyzed on a Bioanalyzer High Sensitivity DNA chip (Agilent). The RNA samples used for library construction had a RIN value range of 7.2 to 9.8, with the median RIN value of all samples being 9.1. Libraries were sequenced as paired end reads with a read length of 100 bp on a HiSeq 3000 (Illumina). ERCC Spike-In Mix (ThermoFisher) was used as an internal control: 2 μL of 1:400 diluted ERCC Spike-In Mix with 500 ng of total RNA input.

For DNA-Seq samples, DNA was extracted using a DNeasy Blood & Tissue Kit (Qiagen, Cat No. 69504). Libraries were generated using Illumina TruSeq DNA PCR-Free Library Prep kit including fragmentation with a Covaris S220 sonic disruptor. Size and purity of the libraries were analyzed on a Bioanalyzer High Sensitivity DNA chip (Agilent). Libraries were sequenced as paired end reads with a read length of 100 bp on an Illumina HiSeq 3000. Six whole genomic DNA samples, comprising three fibropapillomatosis tumor samples and patient-matched non-tumor samples from two juvenile green turtles which had also stranded in Northern Florida, were used for sequencing. These samples were further categorized by tissue type, with one tumor and patient-matched healthy tissue sample each coming from an external, kidney, and lung tissue source (see Supplemental Table 1).

In addition, whole genomic DNA was extracted from 10 plasma samples taken from six individual turtles during the course of their rehabilitation (Supplemental Table 1). Excess blood was taken from blood samples drawn for routine medical care by our veterinarians and plasma was separated from the red blood cells by centrifugation. Only 100 µl of plasma was collected and stored as allowed by permit number MTP-20–236 from the Florida Fish and Wildlife Conservation Commission. DNA was extracted from 60 μl of plasma using a DNeasy Blood & Tissue Kit (Qiagen, Cat No. 69504) and used to produce low input Illumina Fragment libraries using a Low Input Library Prep kit v2 (Clontech Laboratories, Inc., Catalog No. 634899). DNA was fragmented using the Covaris S220 sonic disruptor and libraries were then sequenced as paired end reads with a read length of 100bp on an Illumina HiSeq 3000.

Furthermore, three whole genomic DNA samples were also taken from ground flipper tissue samples from three deceased hatchling turtles, each separate species: a green (*C. mydas*), loggerhead (*Caretta caretta*), and leatherback turtle (*Dermochelys coriacea*), and processed following the methods for tissue as detailed above. The leatherback sequence length was the only sample that differed, with paired end reads of length 150 bp instead of 100 bp.

Finally, one sample of environmental DNA (eDNA) taken from holding tank water from the Whitney Laboratory Sea Turtle Hospital facility of the University of Florida was also used for sequencing. For eDNA sampling, seawater from 5 tanks (4 housing juvenile green sea turtles and 1 housing loggerhead post-hatchling washbacks, 500 ml seawater per tank) was filtered (EMD Millipore PES 0.22um sterivex filter) and DNA was extracted from the filter using a Qiagen DNeasy Blood & Tissue Kit (Qiagen, Cat No. 69504). Libraries were generated using a NEBNext Ultra II DNA Library Prep Kit for Illumina (New England Biolabs, Cat No. E7645), including fragmentation with a Covaris S220 sonic disruptor. Fragment size and purity of the libraries were analyzed on a Bioanalyzer High Sensitivity DNA chip (Agilent). Libraries were sequenced as paired end reads with a read length of 100 bp on an Illumina HiSeq 3000.

### Quality control and read trimming

The software FastQC - https://www.bioinformatics.babraham.ac.uk/projects/fastqc/ - was used to assess data quality. Reads were then trimmed with trim_galore (The Babraham Institute, version 0.5.0) to remove ends with a Phred quality score less than 30, to remove adaptor sequences, and to remove sequences fewer than 25 bp after trimming. For any samples that contained over-represented sequences according to FastQC, the trimmomatic tool ^90^ (version 0.36) was then used to remove these sequences from reads and any sequences less than 25 bp after trimming. The number of raw reads per sample and reads remaining after trimming can be found in Supplemental Table 1.

### Read alignment and read counts

Reads from all samples (RNA-seq, DNA-seq, and eDNA) were first aligned to the ChHV5 genome [GenBank accession number: HQ878327.2]^91^ to determine the level of viral RNA and DNA present in each sample using bowtie2^92^(version 2.3.5.1). The overall alignment rate to the ChHV5 genome was low for both RNA-seq and DNA-seq samples, with < 0.001% of reads aligning on average for all samples (Supplemental Table 1). Reads for all RNA-seq samples were then aligned to the draft genome for *C. mydas* [GenBank assembly accession number: GCA_000344595.1]^93^ using hisat2^94^ (version 2.0.4). The overall alignment rate to the green turtle genome for RNA-seq samples was 82 ± 7% (mean ± SD) (Supplemental Table 1). One sample, an external established growth fibropapillomatosis tumor, had an extremely low alignment rate of 26% to the green turtle genome, and was therefore removed from further analysis. For DNA-seq samples, only tissue, plasma, and the eDNA sample were aligned to the green turtle genome. The overall alignment rates were 91 ± 0.005%, 88 ± 0.02%, and 0.31% (mean ± SD) for tissue samples, plasma samples, and the single eDNA sample, respectively.

Transcript abundance for both ChHV5 virus specific and *C. mydas* specific transcripts was generated using htseq-count^95^ (version 0.6.1p1) with the following parameters: not strand-specific, feature type ‘gene’, and union mode for *C. mydas* specific transcripts, and not strand-specific, feature type ‘gene’, intersection non-empty mode, and a minimum aQual of 0 for ChHV5 virus specific transcripts. Count tables for viral and turtle transcripts were merged for all RNA-seq samples and counts were normalized for gene length and sequencing depth by transcripts per million (TPM) (Supplemental Table 2).

### Differential expression analysis

Prior to differential expression analysis, the raw counts were processed with the RUVseq Bioconductor package ^96^ (version 0.99.1) using the RUVs method to remove low abundance genes, normalize the RNA-seq data, and remove unwanted variation among replicates. PCA plots (see Fig. 1a) were generated using the PtR script in the Trinity toolkit (Haas et al. 2013) both before and after RUVseq normalization. The RUVseq-processed matrix was then used to identify differentially expressed (DE) transcripts using the run_DE_analysis.pl script for the DESeq2 Bioconductor package (Love et al. 2014) and available through the Trinity toolkit (Haas et al. 2013). The run_DE_analysis.pl script was adjusted to also filter out low abundance genes by removing genes with a mean count ≤ 10 across all samples prior to differential expression analysis. The resulting lists of DE genes were sorted and filtered to include only those transcripts with an adjusted p-value of < 0.05 and a log_2_ fold change of > 2 or < −2. A list of upregulated and downregulated transcripts that overlapped from different sample types was generated and used to create area-proportional Venn diagrams of overlap using BioVenn ^97^.

### Pathway analysis and annotation

Gene lists were analyzed for overrepresented pathways, biological functions, and upstream regulators using Ingenuity Pathway Analysis (IPA, Ingenuity Systems, Qiagen). The p-values reported for IPA results were generated by IPA using a right-sided Fisher exact test for over-representation analysis, Benjamini-Hochberg correction for multiple hypothesis testing correction, and a z-score algorithm for upstream analysis; p-values < 0.05 were considered significant. For the systems level analysis, only *C. mydas* DE transcripts that could be annotated to their closest characterized human homolog were included as input.

To better annotate DE transcripts that had turtle-specific gene identifiers [GenBank assembly accession number: GCA_000344595.1], which cannot be used with IPA, the sequence file containing all amino acid sequences for the green turtle genome was re-annotated using PANNZER2 with the –PANZ_FILTER_PERMISSIVE option ^98^. When protein and product descriptions for the annotated DE transcripts agreed between PANNZER2 and the original green turtle genome annotation, the PANNZER2 annotation was used if it provided the name of the closest characterized human homolog instead of a turtle-specific identifier. However, it was often the case that the protein and product descriptions for the annotated DE transcripts were not in agreement, so a random subset of the protein sequences of 11 genes were blasted (blastp, https://blast.ncbi.nlm.nih.gov/Blast.cgi?PAGE=Proteins) against the NCBI non-redundant protein database (nr) to determine which annotation method was most accurate. Since 10 out of the 11 protein sequences tested had the original green turtle genome annotation as the top hit, the genome annotation was used for instances in which the two annotations disagreed. If there was no human homolog available in this case, the genome protein description was checked against the STRING database for human homologs ^99^. Human annotation was used to enable the most comprehensive systems-level analysis, as human genes have been the most extensively annotated and characterized. Out of all of the unique transcripts identified as differentially expressed in all pairwise comparisons, 63% were annotated using the available green turtle genome, 18% were annotated using the PANNZER2 re-annotation of the green turtle genome amino acid sequences, 13% were annotated using the protein description and the STRING database, and 6% of the transcripts either remained unannotated or the annotation was too ambiguous to use in downstream analyses.

### Copy number analysis

To identify potential consistent mutations involved in driving tumorigenesis, copy number variations (CNVs) were identified between paired sets of whole genome DNA sequencing of tumor and healthy tissue each from kidney, lung, and external sources. To assign copy numbers (CNs) to genomic segments of these six genomes, the 140,023 genomic scaffolds of the *C. mydas* draft genome assembly were first combined into a single mega-chromosome. Then, the read-depth based algorithm cn.MOPS ^100^ was used in R to assign copy number, following the methods of Stammnitz et al. (2018)^55^ and using a custom R script produced by the University of Cambridge Transmissible Cancer Group. The script has been made publically available at GitHub (https://github.com/MaximilianStammnitz/turtle-FP-cancer). Samples 27K4H, 27L5H, and liCSVS2 served as normal controls compared to 27K2, 27L1F, and liRRF4 respectively, and read-depths were counted in 5000 bp bins across the combined scaffolds. Segmentation, which determines the length and position of a CNV, was performed using the cn.MOPS algorithm ‘fastseg’ with the referencecn.mops function to join consecutive segments with large or small expected fold changes to make a candidate segment. It then supplies candidate segments that show variations along the mega-chromosome and also across the supplied samples. The custom R script was used to model copy number posterior likelihoods for CN states CN0 through CN128. The locations of the segmented CNVs per scaffold from 27L1F (the only sample with clear copy number gains) were extracted and compared back to the *C. mydas* genome annotation file to compile a list of likely oncogenic genes. Gene lists were analyzed for overrepresented pathways, biological functions, and upstream regulators using IPA as detailed above. For this analysis, only the list of genes was input into IPA due to no available fold-change or p-value for CNVs.

To determine how copy number gains in the genome influence gene expression, RNA-Seq TPM normalized count data was compared between 27L1F (lung tumor with copy number gains) and 27K2F (kidney tumor that was copy number neutral) for each gene found within 27L1F’s higher copy number regions. The expression for each tumor sample was first compared to the expression from its patient-matched healthy tissue sample (27L4H and 27K4H for lung and kidney, respectively) to produce an expression ratio to account for potential tissue specific differences in expression. The expression ratios were then compared between 27-2017-Cm’s lung and kidney tumor samples. If the lung expression ratio was 100% greater than that of the kidney for a specific gene, that gene was considered highly dosage sensitive, in that expression was much higher in the sample with a copy number gain than one that was copy number neutral. If the lung expression ratio was 50% greater than that of the kidney for a specific gene, that gene was considered still slightly dosage sensitive. Any genes that did not fit these criteria, either due to similar expression ratios between the kidney and lung tissues, or due to expression not fitting the expected pattern (i.e. genes being downregulated in the lung tumor relative to healthy lung tissue, which would be expected in genomic regions with copy number losses), were classified as dosage resistant.

### qPCR assays

qPCR assays were conducted on FP tissue samples to quantify viral load within a range of tumor types, as well as eDNA samples to look at viral shedding dynamics within rehabilitation tanks. DNA was extracted from 79 tissue samples from 13 juvenile green turtles which had stranded in Northern Florida as detailed above. DNA was also extracted from 60 tank water samples (20 samplings with three replicates per sampling event). ChHV5 viral load was quantified using TaqMan Fast Advanced Mastermix (ThermoFisher, Cat No. 4444557) according to the manufacturer’s protocol by amplifying the ChHV5 virus-specific DNA polymerase (UL30) gene^25^ (Supplemental Table 4). A species-specific assay was also developed to target the 16S ribosomal RNA gene (rRNA) for Atlantic populations of *C. mydas* to serve as positive controls and to compare the level of viral DNA present against the level of *C. mydas* DNA present within each sample (Supplemental Table 4). This sea turtle species was the primary focus as they are the ones that are most commonly housed at the Whitney Sea Turtle Hospital and most commonly afflicted by FP. A LightCycler490 Instrument II (Roche) was used for amplification and cycling parameters were as follows: 95 °C 10 min, 45 cycles of 95 °C 10 s, 60 °C 20 s, and 72 °C 20 s. All samples were run in triplicate with negative controls. Absolute quantification was utilized in this study. Standard curves using synthetic fragments of the UL30 gene and *C. mydas* 16S rRNA gene (see Supplemental Table 5) were generated to calculate the amount of DNA of these two genes present within each sample (in pg of DNA).

### Histology methodology, embedding, sectioning and staining

Turtle tissue samples surgically removed using a CO_2_ laser and stored in 4% paraformaldehyde at 4°C overnight. Samples washed twice in 1x PBS for 10 minutes; once in Milli-Q H_2_O for 10 minutes; twice in 50% ethanol for 15 minutes; twice in 90% ethanol for 15 minutes; twice in 100% ethanol for 15 minutes. Samples stored in 100% ethanol at 4°C for 3 nights. Samples washed in 100% aniline for 1 hour; 50:50 aniline:methyl salicylate for 1 hour; twice in 100% methyl salicylate for 1.5 hours. Samples stored in 50:50 methyl salicylate:paraffin at 60°C overnight. Samples washed twice in 100% paraffin at 60°C for 3 hours. Samples stored in 100% paraffin overnight. Samples embedded in 100% paraffin and stored at 4°C.

Paraffin blocks sectioned into 6μm ribbons of six on charged Fisherbrand Superfrost Plus microscope slides using an AO Spencer “820” microtome and stored at room temperature

Tissue sections rehydrated by a series of washes: xylene A for 10 minutes; xylene B for 5 minutes; 50:50 xylene:alcohol for 5 minutes; 100% alcohol A for 5 minutes; 100% alcohol B for 5 minutes; 95% alcohol A for 5 minutes; 95% alcohol B for 5 minutes; 80% alcohol for 5 minutes; 70% alcohol for 5 minutes; 50% alcohol for 5 minutes; distilled H_2_O for 5 minutes; 1x PBS. Tissue sections incubated at room temperature for 1.5 hours in 200μl PBS pre-incubation medium (1% normal goat serum + 0.1% bovine albumin serum + 0.1% Tritonx100 + 0.02% sodium azide + PBS). Tissue sections incubated at room temperature overnight in primary antibody medium or control medium (1:100 primary anti β-catenin antibody from rabbit Sigma C2206 + 1:100 primary anti β-actin antibody from mouse Sigma A5441 + PBS pre-incubation medium, or PBS pre-incubation medium only, respectively). Tissue sections washed twice in 1x PBS for 20 minutes. Tissue sections incubated at 37°C for 2 hours in 1:250 FITC GAR (goat anti-rabbit) + TRITC GAM (goat anti-mouse) + PBS pre-incubation medium. Secondary antibodies were from Jackson ImmunoResearch Labaoratires Inc. (West Grove, PA) and were affinity purified and selected for very low cross-reactivity with other animal sources of Ig. Tissue sections washed with 300ml 1x PBS and 2μl Hoechst 33342, trihydrochloride, trihydrate (Life Technologies Corp., Eugene OR) for 10 minutes. Tissue sections washed twice in 1x PBS for 10 minutes. Three drops of 60% glycerol in PBS containing PPD (p-phenylenediamine, 0.3mg/ml) as a fluorescence quench inhibitor were applied to the sections and a cover slip then added to each slide (6 tissue sections per slide). A Leica SP5 confocal microscope was used to visualize and capture images of the fluorescent staining in each tissue section.

For CD3 staining, tissue sections where sent to the University of Florida Veterinary Diagnostic Laboratories core facility, and were stained with rabbit anti-human CD3 ɛ chain antibody clone LN10 (RM-9107-S1 Thermosfisher, Labvision) and an alkaline-phosphatase based red chromogen detection kit. and co-stained with hematoxylin. This CD3 anit-body has previously been validated as also specifically recognizing green sea turtle CD3^56^.

### Retinoic Acid Therapeutic Methodology

Photos with a scale bar were taken of patients undergoing ectopic retinoic acid treatment using an Olympus Tough TG-5, bi-weekly, for the duration of their treatment. This allowed the surface area of each tumor to be analyzed using ImageJ. Direct measurements were also taken bi-weekly using iGaging digital calipers to record the length and width of each tumor. A topical retinoic acid therapeutic (Spear Tretinoin Cream 0.1%) was applied for a 6 - 8 week course depending on the veterinary determination of patient status. Each treated tumor was coupled with a control tumor in the same anatomical location on the opposite side of the body. Tumor length, width, and surface area were analyzed to determine the overall effectiveness of topical retinoic acid treatments for inhibiting FP tumor growth.

### Data availability

The RNA-Seq and DNA-Seq data including raw reads are deposited in NCBI (https://www.ncbi.nlm.nih.gov/) under BioProject ID: PRJNA449022 (https://www.ncbi.nlm.nih.gov/bioproject/PRJNA449022). Code for the copy number analysis has been deposited on Github (https://github.com/MaximilianStammnitz/turtle-FP-cancer/).

## Acknowledgements

Funding was generously provided by The Sea Turtle Conservancy, Florida Sea Turtle Grants Program under project number 17-033R, the Save Our Seas Foundation under project number SOSF 356 and a Welsh Government Sêr Cymru II and the European Union’s Horizon 2020 research and innovation programme under the Marie Skłodowska-Curie grant agreement No. 663830-BU115. This research was also supported by Gumbo Limbo Nature Center, Inc d/b/a Friends of Gumbo Limbo (a 501c3 non-profit organization) through a generous donation through their Graduate Research Grant program. Maximilian Stammnitz is funded by a Gates Cambridge PhD scholarship. Warmest thanks to Nancy Condron, and the veterinary and rehabilitation staff and volunteers of the Sea Turtle Hospital at Whitney Laboratories. Thanks also are due to David Moraga, Yanping Zhang and Mei Zhang of UF’s Interdisciplinary Center for Biotechnology Research Core Facilities and Nicole Stacy and the UF College of Veterinary Medicine Diagnostic Laboratories and Elizabeth Ryan for informative discussions, and Florida Fish and Wildlife Conservation Commission’s Meghan Koperski for valuable assistance with permitting.

---

**Supplemental Figure 1.** Additional gene ontology (GO) term analyses of differentially expressed transcripts. **A - C)** Activation/inhibition z-scores of the top 20 disease-associated GO terms associated with transcripts differentially expressed in different types of fibropapillomatosis tumors (RNA-seq), as detected by IPA, ranked by p-value (calculated by right-tailed Fisher’s Exact Test, with Benjamini-Hochberg correction). **A)** New external FP versus non-tumored skin. A total of 698 of the significant DEs were recognized by IPA and used in the analysis. **B)** Regrowth external FP versus non-tumored skin. A total of 514 of the significant DEs were recognized by IPA and used in the analysis. **C)** Lung FP versus non-tumored lung tissue. A total of 653 of the significant DEs were recognized by IPA and used in the analysis.

**Supplemental Figure 2.** Additional fibropapillomatosis transcriptomic comparisons. **A)** Overlap of transcripts from RNA-seq data significantly differentially expressed (DE) (as called by DESeq2) in fibropapillomatosis between either kidney FP compared to established external FP, or lung FP compared to established external FP. Area-proportional diagrams were generated using BioVenn (http://www.biovenn.nl/). Transcripts were considered significant if passing the following cut-offs: adjusted p-value of < 0.05 and log_2_ fold change of > 2 and < −2. **B, C)** Activation/inhibition z-scores of the top 20 disease-associated gene ontology (GO) terms associated with transcripts differentially expressed in different types of fibropapillomatosis tumors (RNA-seq), as detected by IPA, ranked by p-value (calculated by right-tailed Fisher’s Exact Test, with Benjamini-Hochberg correction). **B)** Kidney FP versus established external FP tumors. **C)** Lung FP versus established external FP tumors. **D)** Activation z-scores of the Interferon gamma (IFNG) inferred transcriptional regulator (ITR) associated with transcripts differentially expressed across the five different types of fibropapillomatosis tumors, when compared to their respective non-tumor tissue sources.

**Supplemental Figure 3. A)** Time-course of relative tumor growth profiles of retinoic acid (RA, Tretinoin cream 0.1%) treated and untreated fibropapillomatosis tumors. Profiles for eight tumors across three individual *C. mydas* patients are shown. Duration of treatment was under veterinary determination. Patient ‘name’ abbreviations, Ferd., ‘Ferdinand’ (07-2018-Cm), Eins., ‘Einstein’ (28-2018-Cm) and ‘Lilac’ (25-2018-Cm). Tumor growth is relative to the size of each individual tumor before treatment, i.e. Day 1.

**Supplemental Figure 4.** Additional fibropapillomatosis tumor immunohistochemistry. **A)** Anti-body based immunohistochemistry of established external, regrowth external, regrowth eye external and kidney internal tumor tissue. Tissue sections are stained for β-catenin (anti-β-catenin antibody) and counter stained with Hoechst 33342 to visualize nuclei and Anti β-actin. Selected cells with nuclear (activated) β -catenin staining are indicated by white arrows.

**Supplemental Figure 5.** Additional fibropapillomatosis inferred transcriptional regulator (ITR) interaction networks. **A, B)** Interaction networks of the top 200 ITRs of established **A)** external tumors and of **B)** lung tumors. Networks were generated by String^99^ (https://string-db.org/).

**Supplemental Figure 6.** Additional fibropapillomatosis inferred transcriptional regulator (ITR) network-based functional module discovery. **A-C)** Network-based functional module discovery of the top 200 ranked ITRs (called by IPA) of **A)** kidney tumors **B)** new growth external tumors and **C)** regrowth external tumors. Networks were generated using HumanBase (https://hb.flatironinstitute.org/). **D)** Activation z-scores of the ‘Quantity of Metal’ gene ontology (GO) term associated with transcripts differentially expressed in different types of fibropapillomatosis tumors (kidney FP, lung FP, external FP) when compared to their respective non-tumored tissue sources, as detected by IPA. Note: the kidney FP disease-associated GO analysis did not have the ‘Quantity of Metal’ GO term called when compared to healthy kidney tissue.

**Supplemental Figure 7.** Additional copy number variation (CNV) data. **A)** Genome-wide copy number variation (CNV) analysis of whole genome sequencing of patient-matched tumor and non-tumor tissue (top: kidney tumor compared to healthy kidney from patient 27-2017-Cm; bottom: early external tumor compared to non-tumor skin from 25-2018-Cm, ‘Lilac’). Copy number variation analysis was carried out using the cn.MOPS package^100^ in R. Tumored tissue was compared to non-tumored tissue from the same tissue source and same patient for each comparison. Each point shows a genomic segment 5000 bp long and its associated copy number within the tumor relative to its patient-matched non-tumor tissue, with a copy number of 2 being the null hypothesis for a diploid organism. Red data points indicate where the segmentation algorithm called a potential significant CNV. Both samples shown above are examples of a copy number neutral state. **B**) Network-based functional module discovery of the genes harboring CNV (gains) in 27-2017-Cm’s lung fibropapillomatosis tumor. Network was generated using HumanBase (https://hb.flatironinstitute.org/).

**Supplemental Figure 8.** Additional gene level ChHV5 transcriptomics analysis. **A)** Bar graph showing level of ChHV5 viral transcript expression of every gene in the ChHV5 genome, averaged by tissue type. Reads were first aligned from RNA-seq samples to the ChHV5 genome^91^ via bowtie2 and read counts generated per viral gene using htseq-count. Reads were normalized by gene length and sequencing depth to transcripts per million (TPM) and averaged across samples for the following groups: non-tumor tissue n = 20, external fibropapillomatosis tumors n = 56, lung fibropapillomatosis tumors n = 7, and kidney fibropapillomatosis tumors n = 6.

**Supplemental Figure 9.** ChHV5 transcript levels compared with rehabilitation outcome. **A)** Bar graph showing total level of ChHV5 RNA among samples by individual patient, as assessed by RNA-seq. Reads were normalized by sequencing depth to number of viral reads per 10 million total reads per sample. Bars show individual samples per patient. **B)** Dot plot of the number of ChHV5 reads (RNA-Seq) per 10 million total reads between patients based on outcome (released vs died in care/humanely euthanized). Line denotes mean, error bars denote standard error. Significant difference in averages between the two outcomes was determined by a Mann-Whitney U Test and is denoted by an asterisk (*). Per outcome: released = 7 turtles; died/euthanized = 5 turtles.

**Supplemental Figure 10.** Additional gene level ChHV5 genomic (DNA-seq) analysis. **A)** Bar graph showing level of ChHV5 viral DNA reads of every gene in the ChHV5 genome. For the three non-tumor samples reads per gene were average. Reads were first aligned from DNA-seq samples to the ChHV5 genome^91^ via bowtie2 and read counts generated using htseq-count. Reads were normalized by gene length and sequencing depth to transcripts per million (TPM). Per sample type: nontumor tissue n = 3, external FP tumor n = 1, lung FP tumor n = 1, and kidney FP tumor n = 1.

