## Supplementary figures and images for "Mutational, transcriptional and viral shedding dynamics of the marine turtle fibropapillomatosis tumor epizootic"

### Supplemental Figure 1

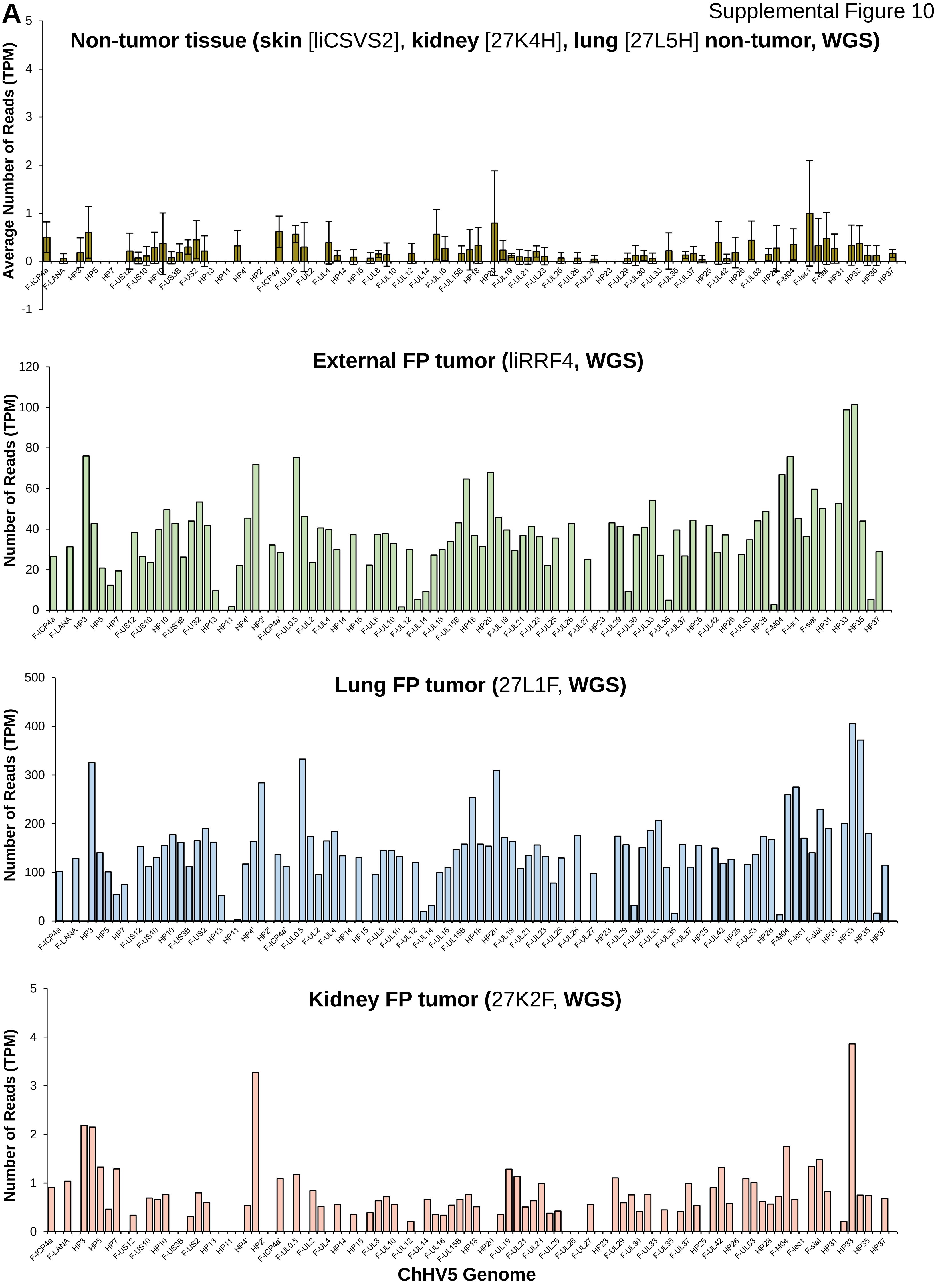

### Supplemental Figure 1.

Supplemental Fig. 1

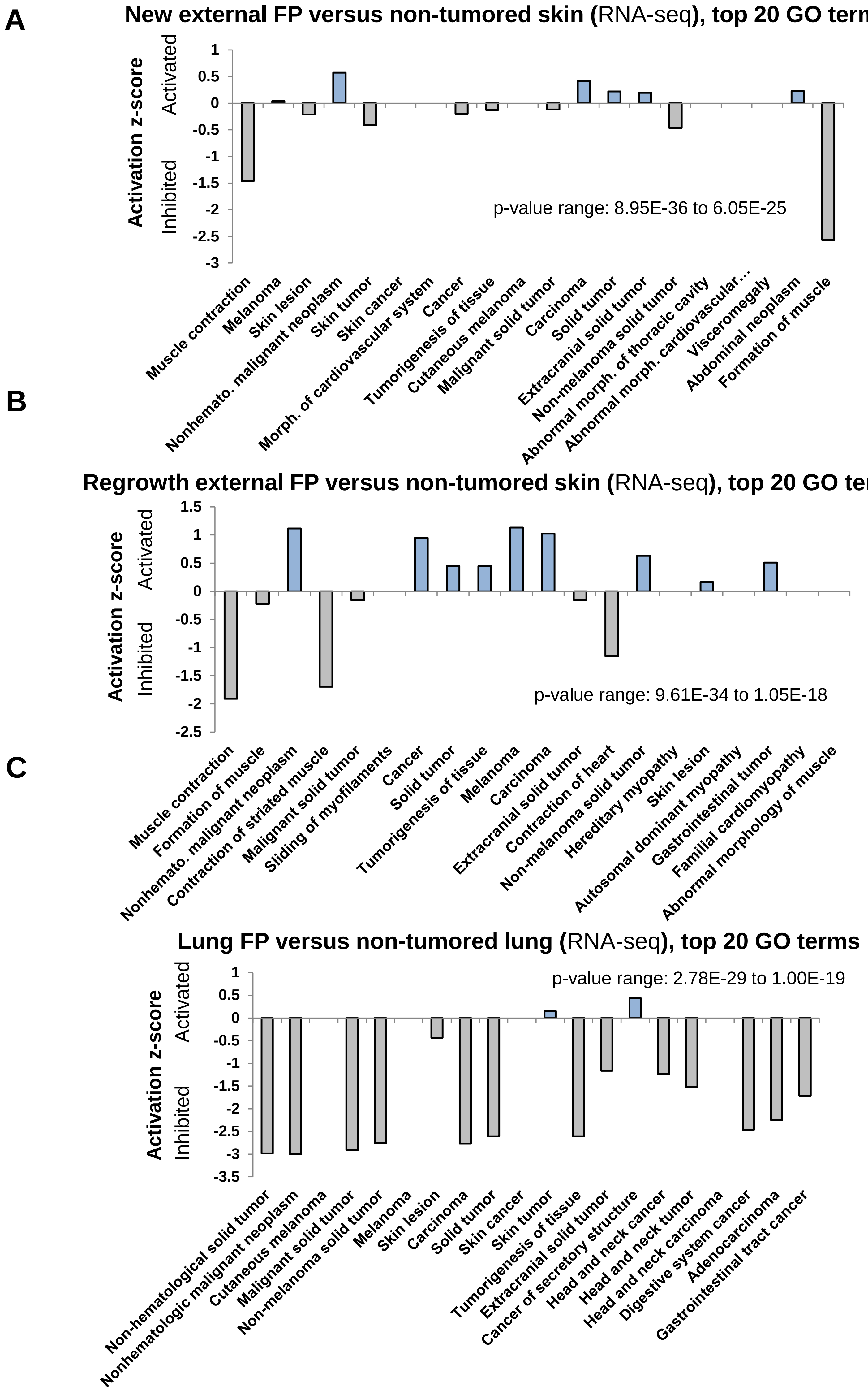

### Supplemental Figure 2.

Supplemental Figure 2

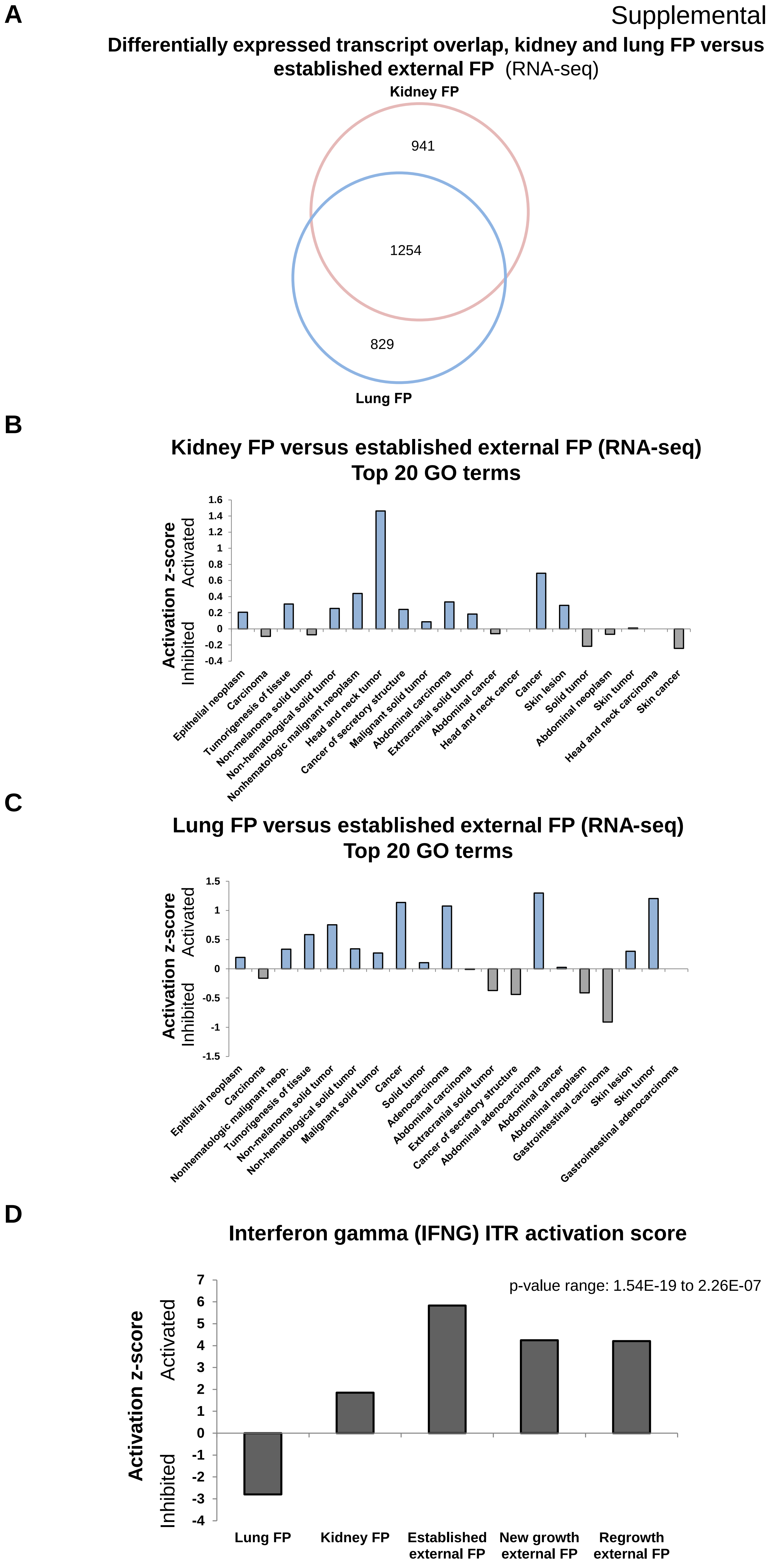

### Supplemental Figure 3.

**A** Supplemental Figure 3

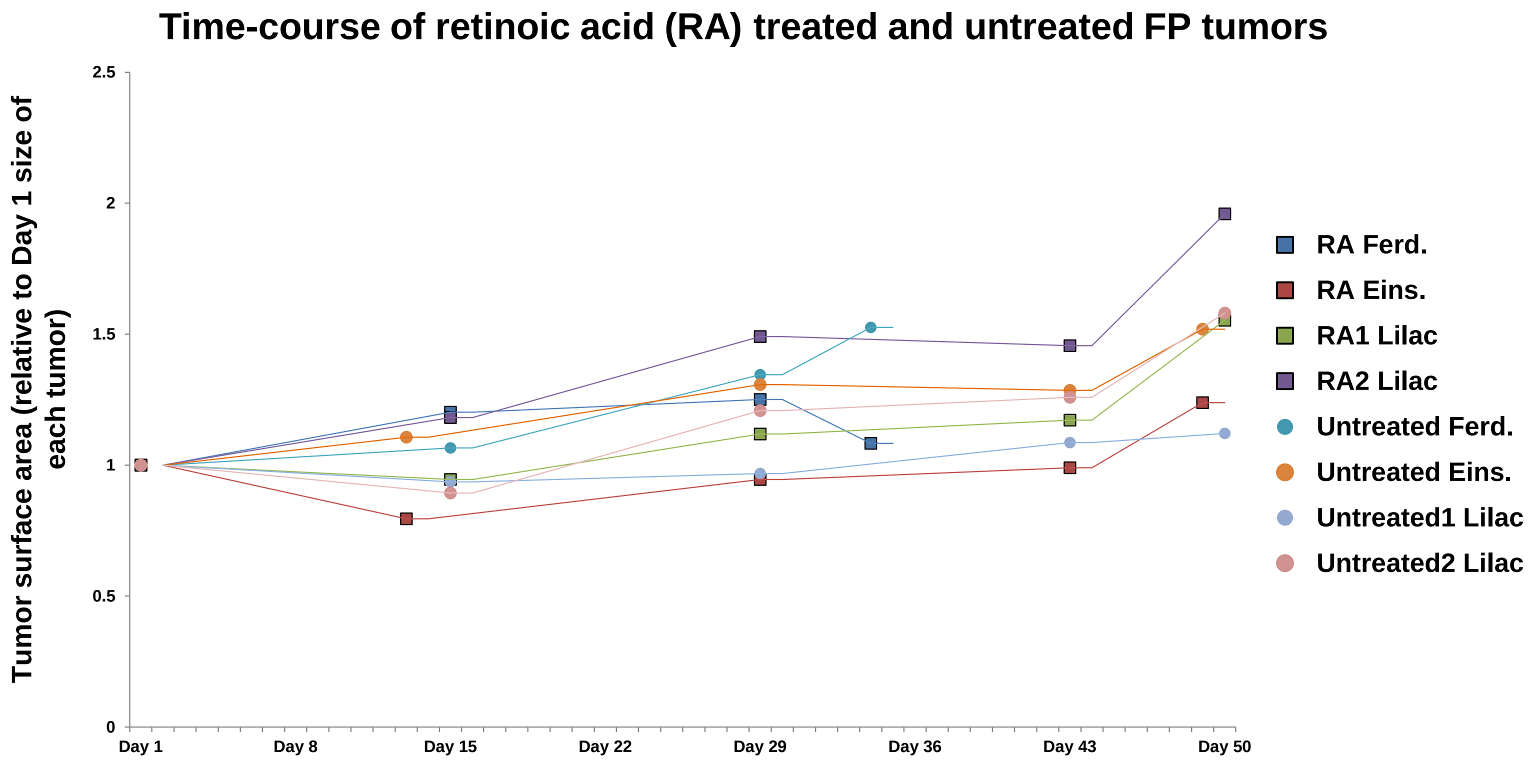

### Supplemental Figure 4.

Supplemental Figure 4

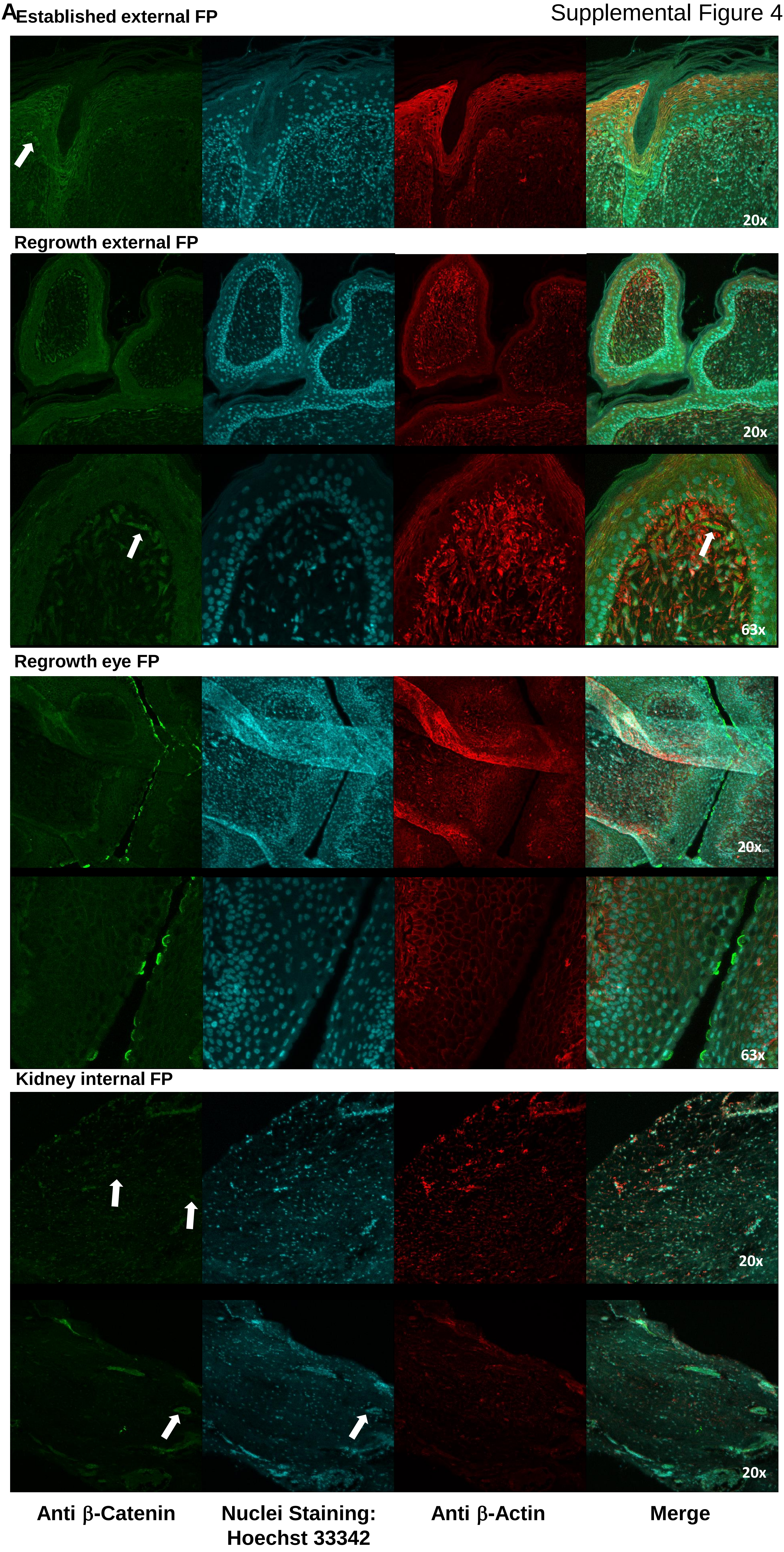

### Supplemental Figure 8.

A

Supplemental Figure 8

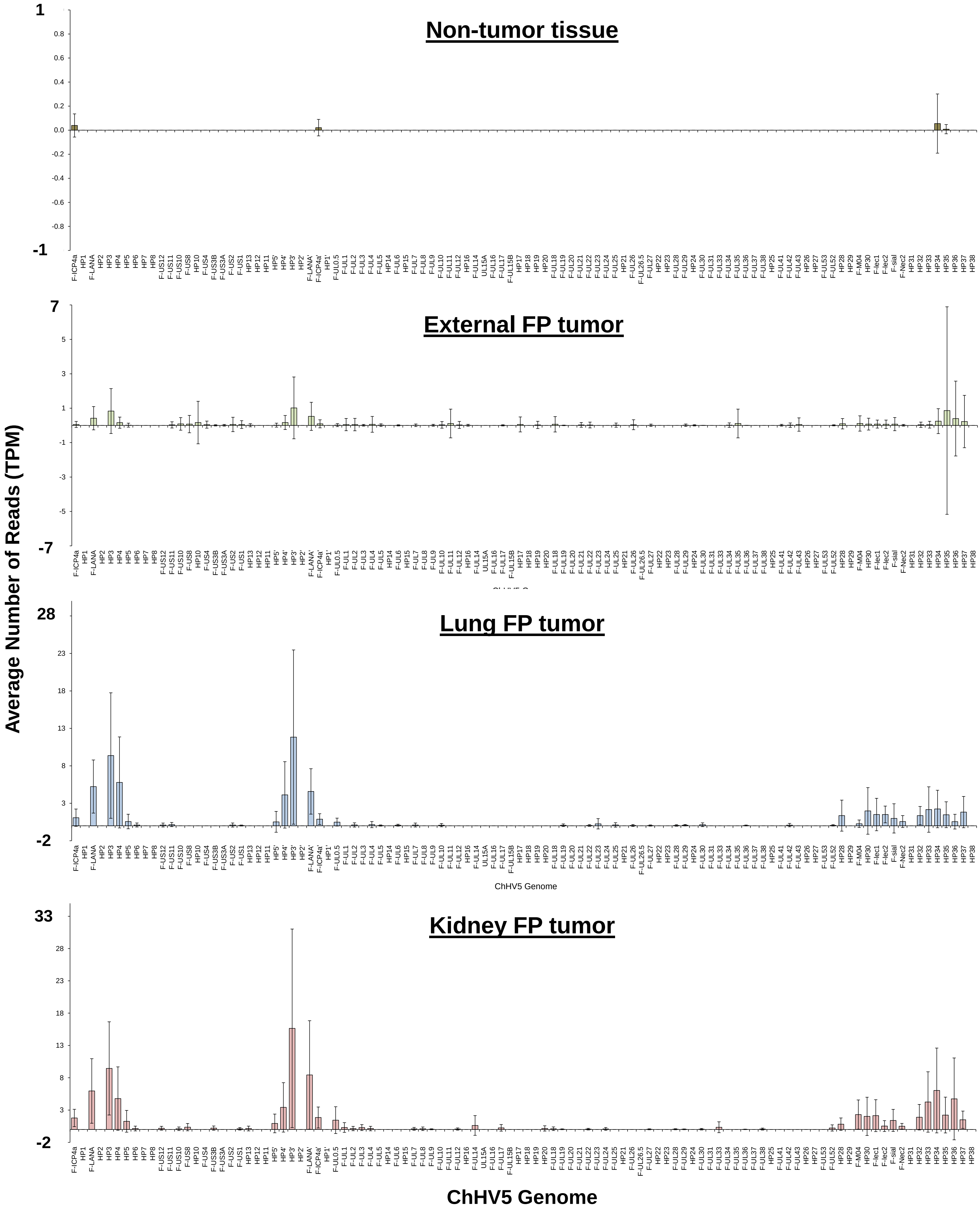
