## Supplemental Figure 5. for "Mutational, transcriptional and viral shedding dynamics of the marine turtle fibropapillomatosis tumor epizootic"

**A**

### Established external FP versus non-tumored skin (RNA-seq)
### Top 200 ITRs

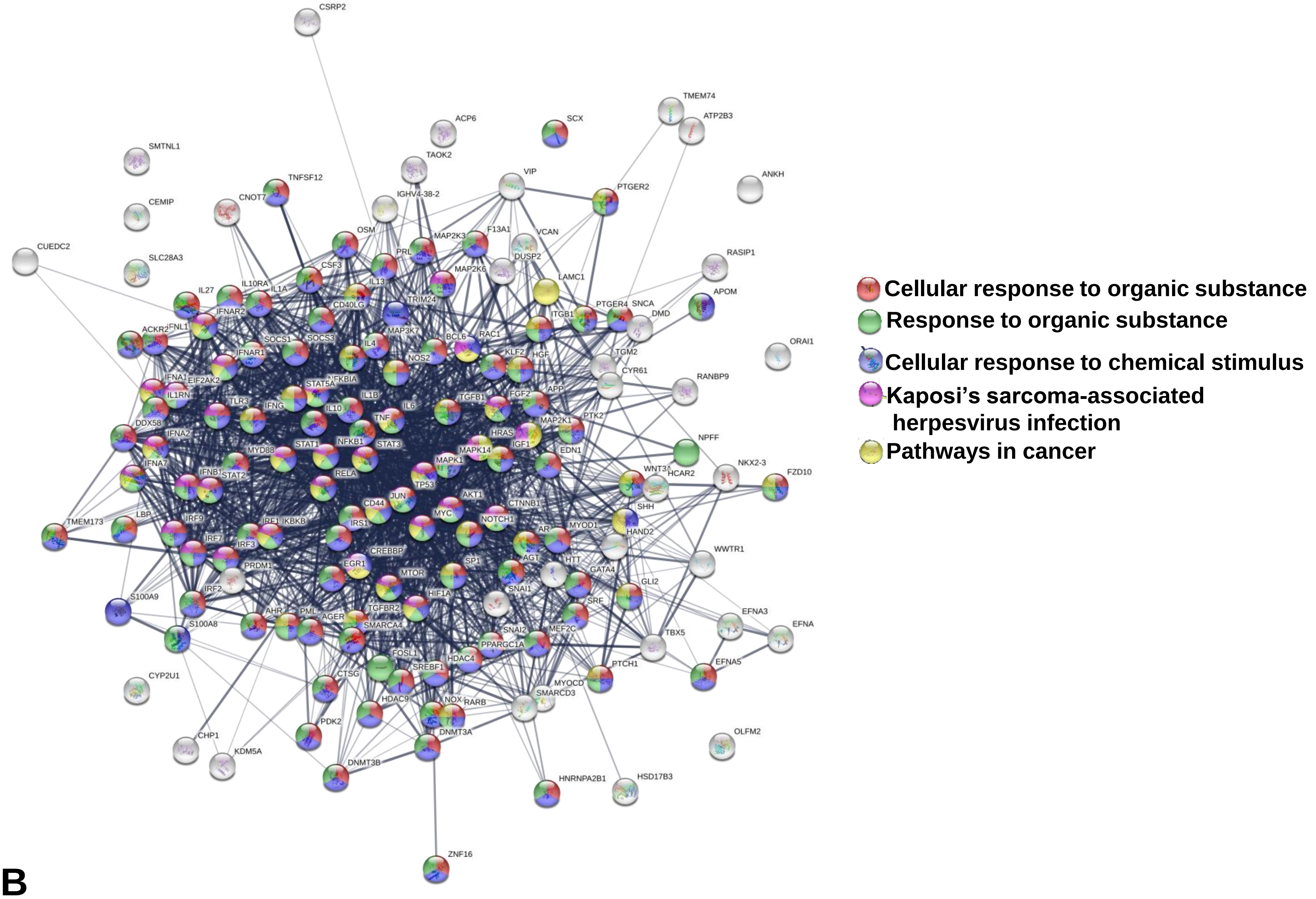


**B**

### Lung FP versus non-tumored skin (RNA-seq)
### Top 200 ITRs

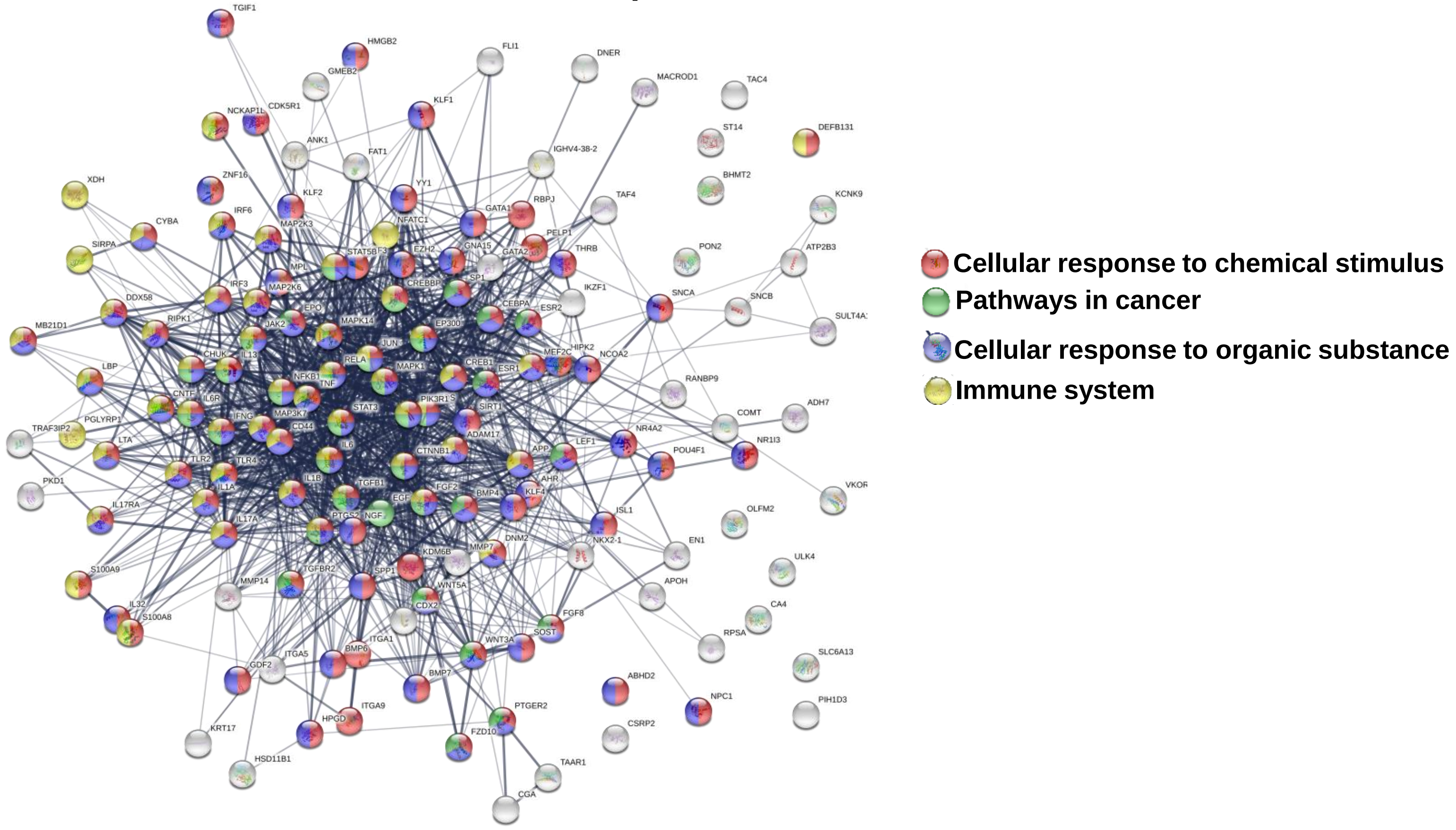
