## Supplemental Figure 6. for "Mutational, transcriptional and viral shedding dynamics of the marine turtle fibropapillomatosis tumor epizootic"

**A**

### Kidney FP versus non-tumored kidney tissue top 200 ITR network (RNA-seq)

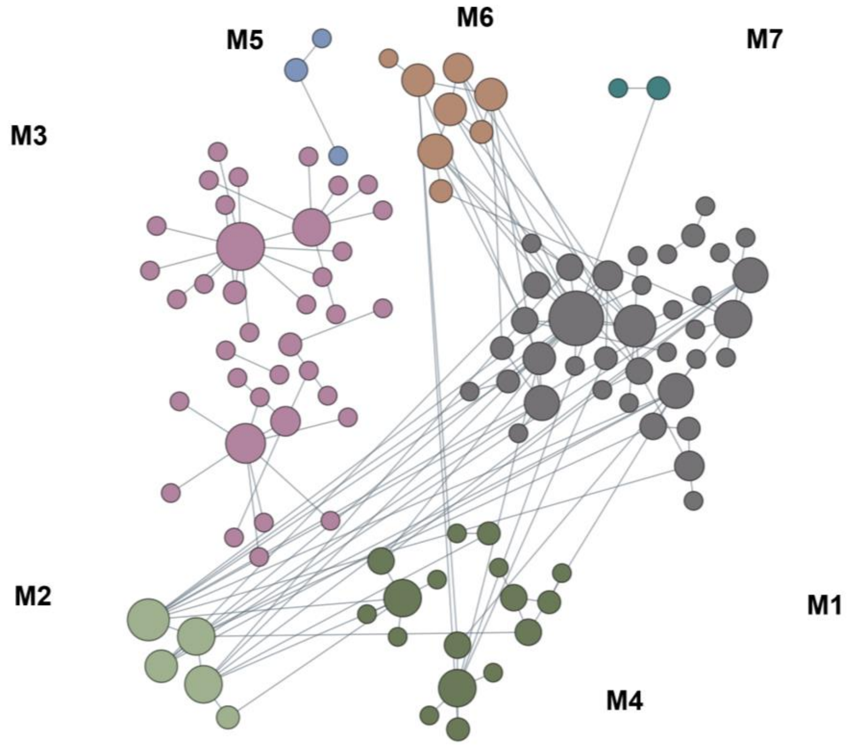


| | Kidney Top term | Q value |
|---|---|---|
| M1 | Response to growth factor | <1e-04 |
| M2 | Cellular response to interleukin 6 | <1e-04 |
| M3 | Epithelium development, embryo development | <1e-04 |
| M4 | Positive regulation of protein kinase activity | <1e-04 |
| M5 | Positive regulation of protein chemokine secretion | <1e-04 |
| M6 | Myeloid leukocyte development | <1e-04 |
| M7 | Cellular response to growth factor | 0.0010 |

**B**

### New growth external FP versus non-tumored skin top 200 ITR network (RNA-seq)

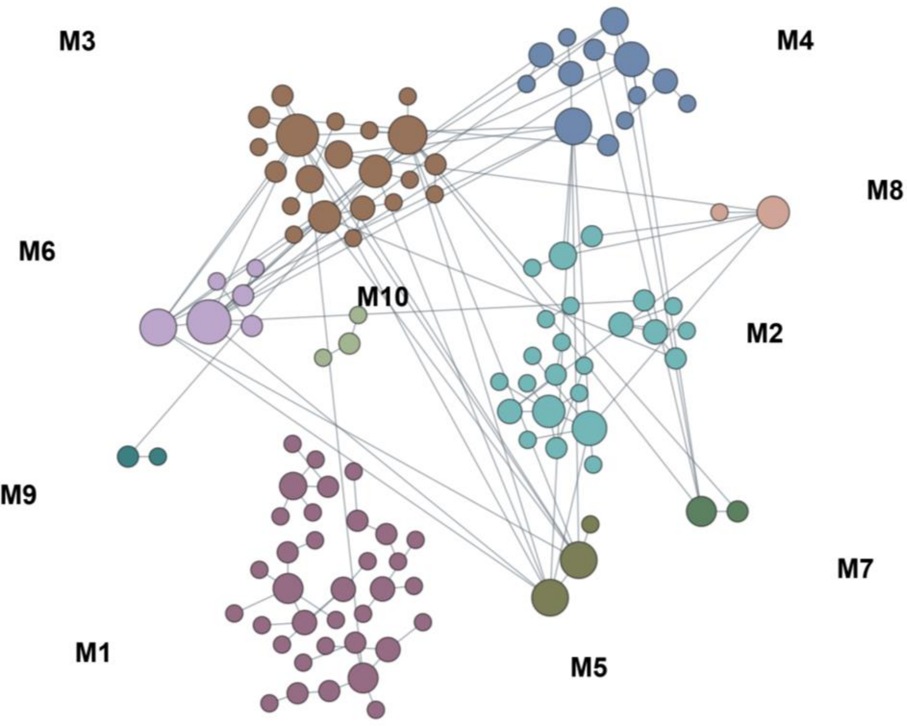


| | Top term | Q value |
|---|---|---|
| M1 | Regulation of JAK-STAT cascade | <1e-04 |
| M2 | Regulation of DNA binding transcription and viral transcription | <1e-04 |
| M3 | Positive regulation of cytokine production | <1e-04 |
| M4 | Positive regulation of pri-miRNA transcription | <1e-04 |
| M5 | Hyaluronan metabolic process | <1e-04 |
| M6 | Positive regulation of pri-miRNA transcription | 0.0001 |
| M7 | Cellular response to reactive oxygen species | 0.0002 |
| M8 | Cell cycle arrest, negative regulation of cell growth, Ras protein signal transduction | 0.0002 |
| M9 | Positive regulation of protein/peptide secretion | 0.0004 |
| M10 | Regulation of leukocyte cell-cell adhesion | 0.0007 |

**C**

### Regrowth external FP versus non-tumored skin top 200 ITR network (RNA-seq)

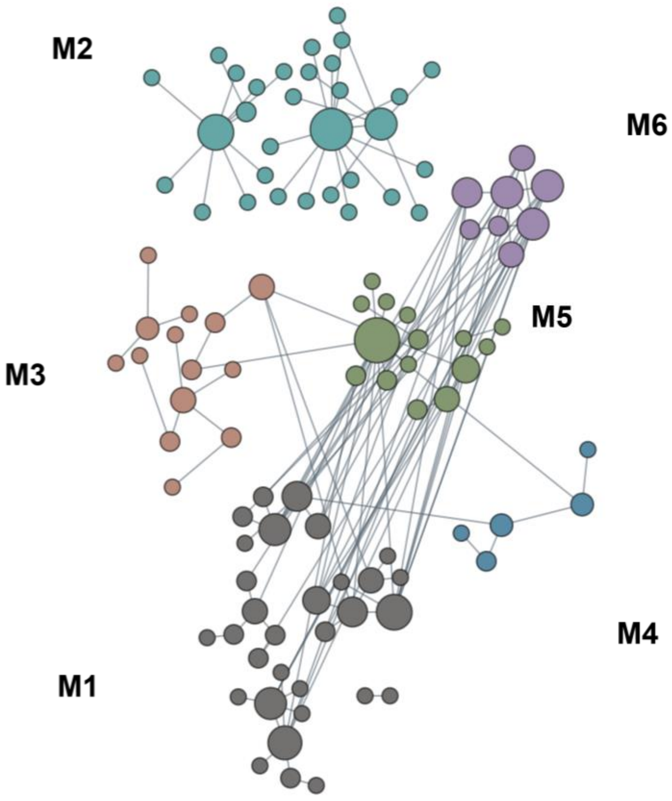


| | Top term | Q value |
|---|---|---|
| M1 | Cellular response to cytokine stimulus | <1e-04 |
| M2 | Cytokine production involved in immune response | <1e-04 |
| M3 | Positive regulation of cell mobility (EMT) | <1e-04 |
| M4 | Cellular response to peptide hormone | <1e-04 |
| M5 | DNA methylation on cytosine | <1e-04 |
| M6 | Regulation of response to external stimuli (chemotaxis) | 0.0001 |

**D**

### Quantity of Metal GO term

(RNA-seq)

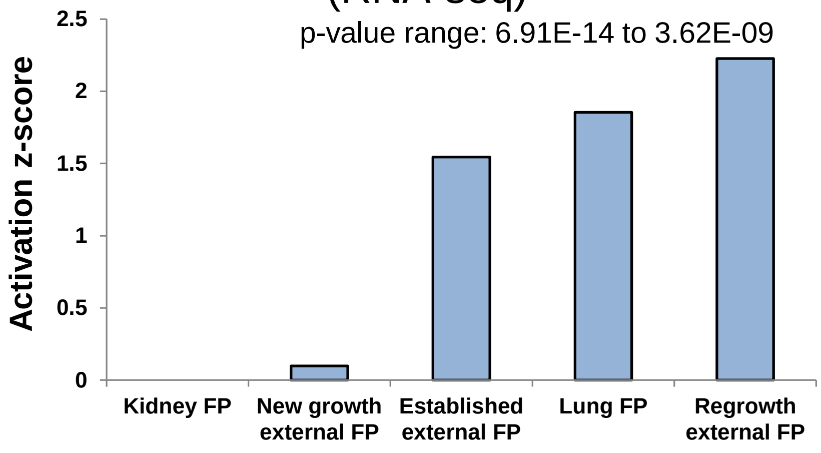
