## Supplemental Figure 7. for "Mutational, transcriptional and viral shedding dynamics of the marine turtle fibropapillomatosis tumor epizootic"

**A**

**Genome-wide copy number variation (CNV)**

Patient 27-2017-Cm Kidney FP Vs non-tumor kidney tissue (WGS, DNA-seq)

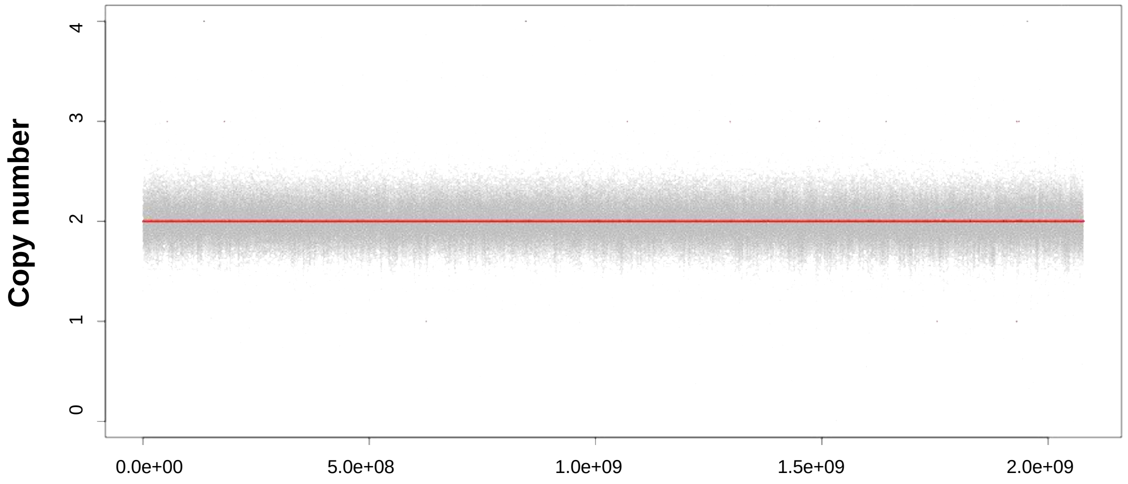


Patient Lilac (25-2018-Cm) early External FP Vs non-tumor skin tissue (WGS, DNA-seq)

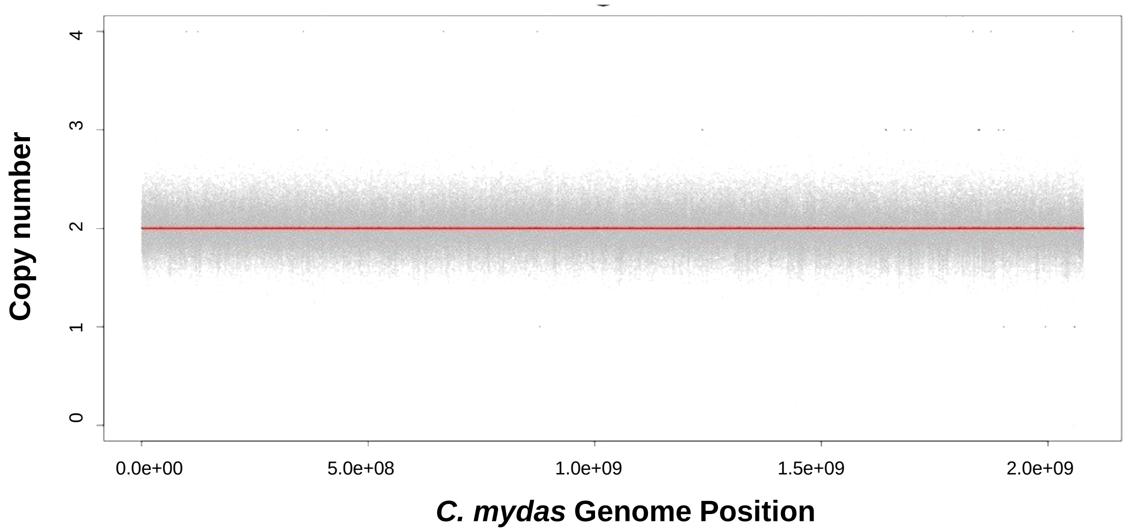


**B**

**Network of genes with CNV in 27-2017-Cm Lung FP tumor**

(DNA-seq, 27-2017-Cm Lung FP Vs 27-2017-Cm non-tumor lung tissue)

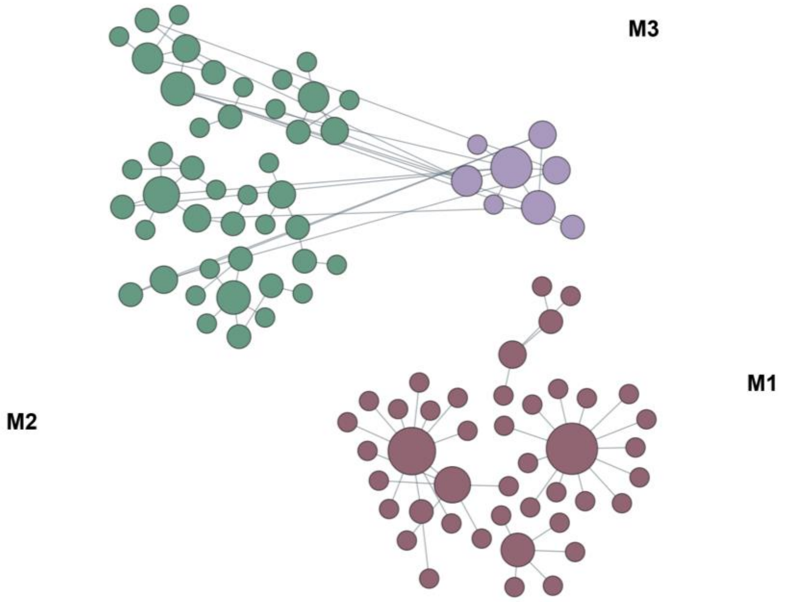


| | Top term | Q value | Genes (ITRs) | Terms |
|---|---|---|---|---|
| **M1** | Embryonic morphogenesis | 0.0002 | 43 | 49 |
| **M2** | Ribonucleoprotein complex assembly/negative regulation of mitotic cell cycle phase transition/DNA replication checkpoint | 0.0004 | 44 | 76 |
| **M3** | Head development | 0.0057 | 8 | 1 |
