## Supplemental Figure 9. for "Mutational, transcriptional and viral shedding dynamics of the marine turtle fibropapillomatosis tumor epizootic"

**A**

### ChHV5 viral reads across individual samples, arranged by turtle of origin (RNA-seq)

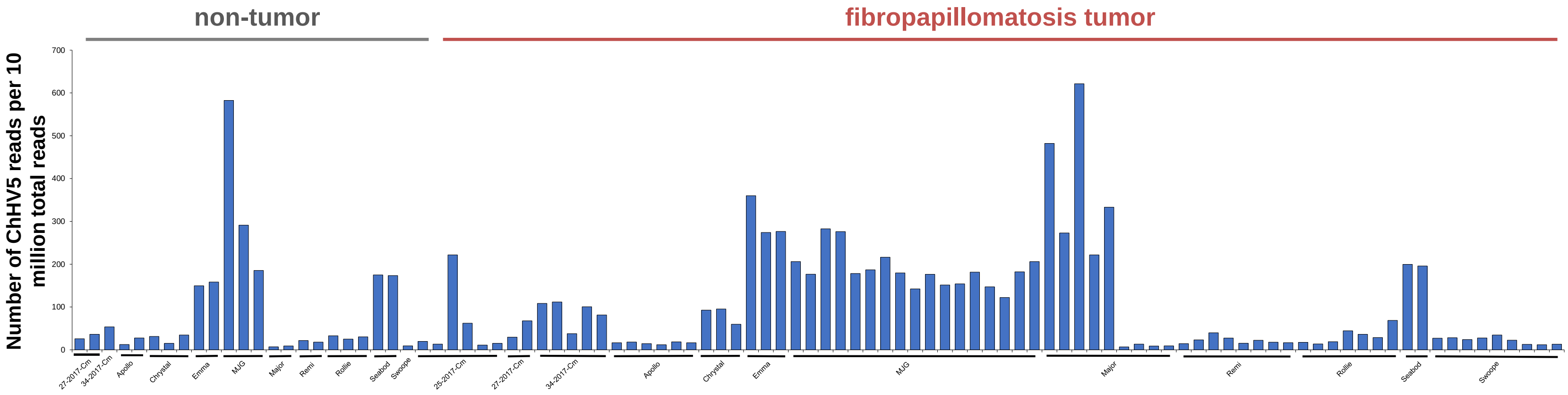


**B**

### ChHV5 viral reads across individual samples, arranged by patient outcome (RNA-seq)

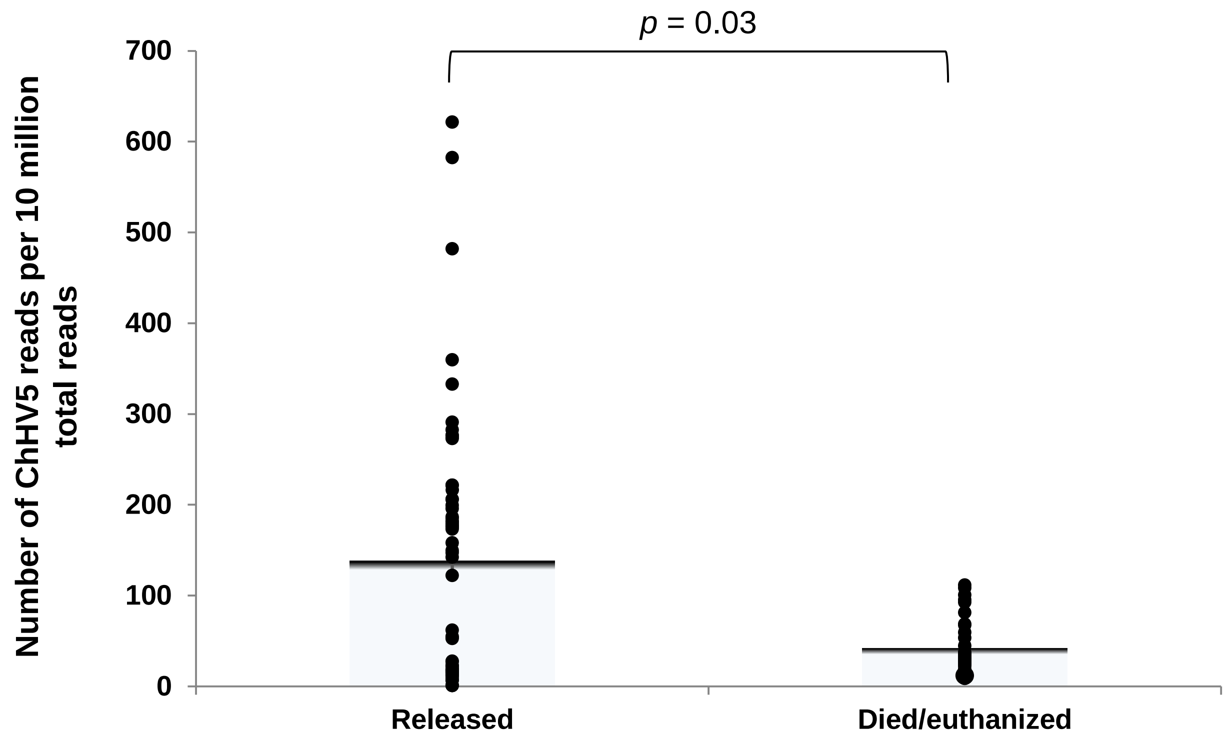
